# Amycomicin: a potent and specific antibiotic discovered with a targeted interaction screen

**DOI:** 10.1101/317362

**Authors:** Gleb Pishchany, Emily Mevers, Sula Ndousse-Fetter, Dennis J. Horvath, Camila R. Paludo, Eduardo A. Silva-Junior, Sergey Koren, Eric P. Skaar, Jon Clardy, Roberto Kolter

**Author notes:** These authors contributed equally (Gleb Pishchany, Emily Mevers).

## Abstract

The rapid emergence of antibiotic-resistant pathogenic bacteria has accelerated the search for new antibiotics. Many clinically used antibacterials were discovered through culturing a single microbial species under nutrient-rich conditions, but in the environment, bacteria constantly encounter poor nutrient conditions and interact with neighboring microbial species. In an effort to recapitulate this environment we generated a nine-strain *Actinomycete* community and used 16S rDNA sequencing to deconvolute the stochastic production of antimicrobial activity that was not observed from any of the axenic cultures. We subsequently simplified the community to just two strains and identified *Amycolatopsis* sp. AA4 as the producing strain and *Streptomyces coelicolor* M145 as an inducing strain. Bioassay-guided isolation identified amycomicin, a highly modified fatty acid containing an epoxide isonitrile warhead as a potent and specific inhibitor of *Staphylococcus aureus.* Amycomicin targets an essential enzyme in fatty acid biosynthesis (FabH) and reduces *S. aureus* infection in a mouse skin infection model. The discovery of amycomicin demonstrates the utility of screening complex communities against specific targets to discover small-molecule antibiotics.

**Significance:** Bacteria, especially actinomycetes, produce the majority of our clinically useful small-molecule antibiotics. Genomic analyses of antibiotic-producing strains indicate that earlier discovery efforts found only a fraction of the likely antibiotic candidates. In an effort to uncover these previously missed candidates we developed an approach that utilizes the ability of microbial communities to produce antibiotics that are not produced by any single member in isolation. Successful communities were established and deconvoluted to identify both producers and inducers of antibiotic activity. One inducer-producer pair made amycomicin, a potent and specific antibiotic against *Staphylococcus aureus*, an important human pathogen. Amycomicin targets fatty acid biosynthesis and exhibits *in vivo* efficacy against skin infections in a mouse model.

## Introduction

Researchers have responded to the increasing incidence of antibiotic-resistant bacterial infections with attempts to discover new small molecule antibiotics (1, 2). Many of these attempts apply new techniques to traditional sources, including the environmental bacteria that have provided the suite of antibiotics currently being threatened by resistance (3). This return to traditional sources began with the realization that earlier explorations had captured only a small fraction of the biosynthetic potential seen in bacterial genomes (2, 4). These missing molecules, frequently called ‘cryptic’ metabolites, can be inferred by identifying their biosynthetic genes but have not yet been confirmed in the laboratory (5). Previous studies may have missed these molecules because they are produced on an ‘as-needed’ basis and culturing a single strain in rich media may not create the requisite need.

Bacteria sense and respond to the world around them with small molecules, and adding non-kin microbes, environmental, or host factors during culturing can induce production of cryptic metabolites (6-15). We have developed a discovery approach in which metabolite expression is induced through co-culturing interactions in a complex community and simultaneously linked to a therapeutically relevant target assay, an approach we call a targeted-interaction screen.

For the interacting organisms, we used a collection of nine previously studied *Actinomycete* strains. These bacteria have been a prolific source of antibiotics and their social lifestyles suggested that they may produce numerous small molecules in response to neighboring microbes (3, 9, 10, 16). A dilute complex medium was used for culturing because previous studies indicated that such conditions are conducive to the growth of co-existing strains as well as interspecific induction of antibiosis (10, 17, 18). For the target, we selected the important human pathogen, *Staphylococcus aureus*. After an initial screen to reduce community complexity, we focused on a strong response in which *Streptomyces coelicolor* M145 produces a signal that causes *Amycolatopsis* sp. AA4 to produce an antibiotic that potently (MIC ~30 nM) and specifically kills *S. aureus* in both *in vitro* and *in vivo* assays. Further analysis led to the identity of the antibiotic, which we named amycomicin (AMY). AMY is an unusual fatty acid-based antibiotic that has been highly functionalized, including incorporation of a ketone, a hydroxyl, an isonitrile-epoxide, and an olefin. We also identified AMY’s biosynthetic gene cluster in AA4, which contains a recently identified isonitrile synthase gene (19), and AMY’s target as FabH, an enzyme in the fatty acid biosynthesis pathway. Finally, AMY is effective in a murine skin infection model of *S. aureus.*

## Results and Discussion

### Inter-strain interactions induce production of antimicrobial activity

In *Actinomycetes* antibiotic production is linked to developmental stage, so we used solid phase culturing in transwell plates allowing continuous monitoring without disrupting the community (Fig. 1A) (16). A transwell sealed by a permeable membrane, is filled with a 0.75% agarose solution. Once the agarose solidifies, the transwell is partially submerged into liquid medium within a larger well. *Actinomycete* spores are then inoculated on the agarose surface. The nutrients present in the liquid medium diffuse through the membrane and the agarose plug, allowing the bacteria to grow. As the bacterial community proliferates, secreted metabolites diffuse through the agarose into the liquid medium. The conditioned liquid medium can then be periodically tested for antibiotic activity and/or exchanged to provide fresh nutrients.

To capture multiple interactions, we began with a nine-strain community of *Actinomycetes.* Spores were inoculated onto the 200 μL agarose plugs of the transwell system singly or as a mix of all nine species. One milliliter of dilute complex medium (0.1X PTSYM, See Methods), was added to the bottom wells. To assess secreted antimicrobial activity, 10 μL samples were periodically removed from the bottom well and spotted on agar plates freshly seeded with a lawn of *S. aureus* (Fig. 1A). Medium was replaced after three weeks.

**Fig. 1.**
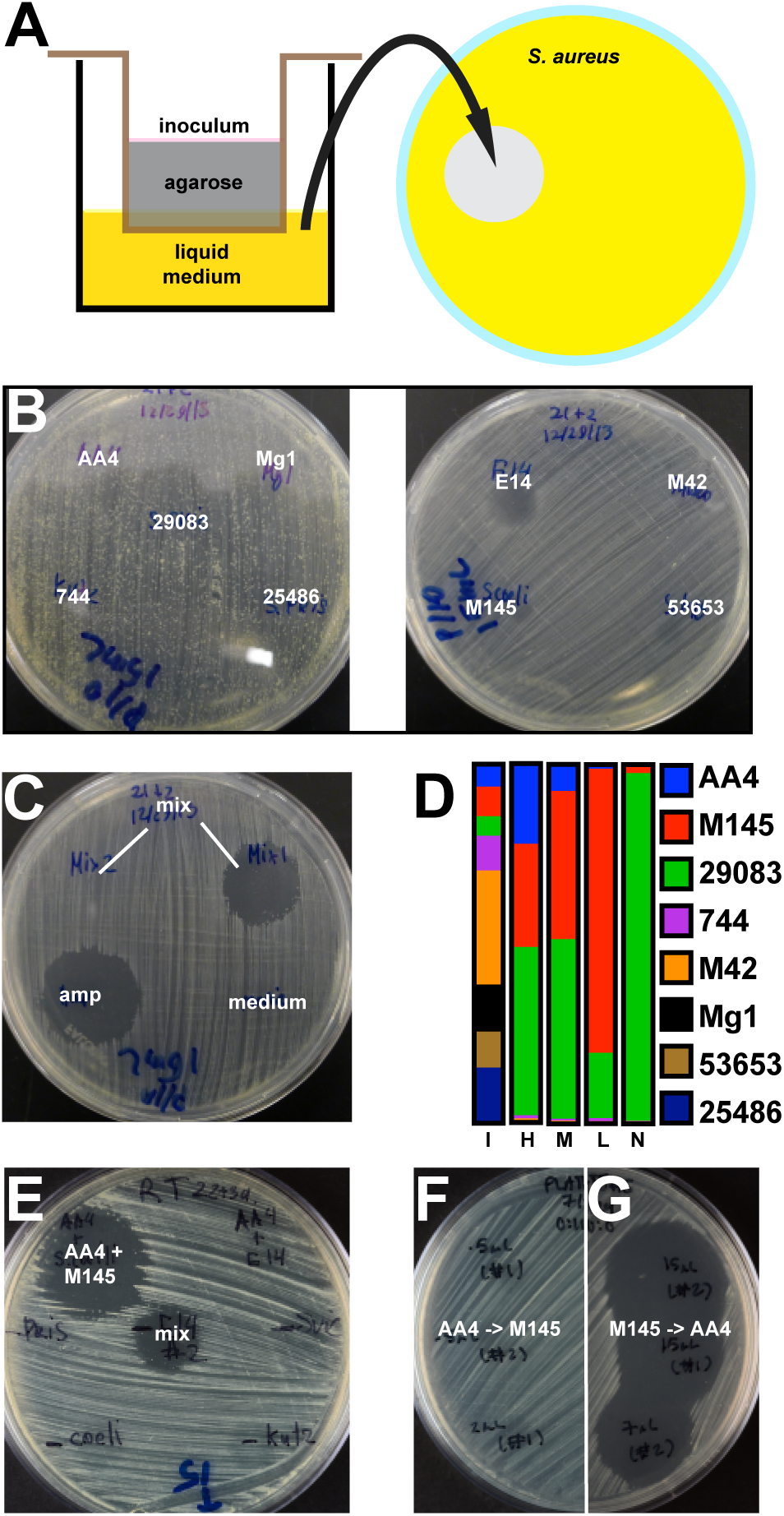
Antimicrobial activity induced by interspecies interactions in *Actinomycete* communities. (*A*) Microbial strains or communities were inoculated onto an agarose plug that was supported by a permeable membrane within a transwell. The conditioned liquid medium was periodically sampled for antibiotic (see Materials and Methods for details). (*B*) Individual strains do not produce strong antimicrobial activity against *S. aureus. Amycolatopsis* sp. AA4 (AA4), *Streptomyces* sp. Mg1 (Mg1), *Streptomyces sviceus* ATCC 29083 (29083), *Kutzneria* sp. 744 (744), *Streptomyces pristinaespiralis* ATCC 25486 (25486), *Streptomyces* sp. E14 (E14), *Micromonospora* sp. M42 (M42), *Streptomyces coelicolor* A3(2) M145 (M145), *Streptomyces hygroscopicus* ATCC 53653 (53653) (*C*). A community of all nine strains (mix) sometimes produced and sometimes did not produce antimicrobial activity. Ampicillin (amp) and fresh medium (medium) were used as positive and negative controls, respectively. (*D*) Relative abundance of each strain in the microbial communities, as determined by 16S rDNA sequencing, at the time of inoculation (I), and at the time when antibiotic was produced in communities that produced relatively high (H), medium (M), low (L) or no activity (N). (*E*) Zone of inhibition produced by a bipartite community of AA4 and M145 (AA4 + M145). Other combinations tested (AA4 + E14…) did not produce inhibitory activity. A sample from the mix was placed in the middle as a control. (*F* and *G*) Zones of growth inhibition produced by medium that was sequentially conditioned by AA4 and then M145 (*F*) or by M145 and then AA4 (*G*).

After 1-2 additional days, we sometimes detected the production of a strong antimicrobial activity against *S. aureus.* While none of the individual strains produced strong antibiosis (Fig. 1B), cultures that started with approximately equal numbers of all nine strains occasionally produced a strong activity (Fig. 1C). This variability was attributed to the stochasticity of complex microbial communities (20, 21).

We used 16S rDNA sequencing to quantify each strain’s relative abundance at the time of antibiotic production (Fig. 1D). At the time antimicrobial activity was detected, the communities were dominated by three of the original nine strains: *Amycolatopsis* sp. AA4 (henceforth AA4), *Streptomyces coelicolor* A3(2)/M145 (henceforth M145), and *Streptomyces sviceus* ATCC 29083. Most importantly, the proportion of AA4 present correlated loosely with activity. For example, the community producing the most activity (column H in Fig. 1D) contained 22% AA4, while a community with no detectable activity (column N in Fig. 1D) contained only 0.11% AA4. Ultimately, we found robust antibiotic productions in co-cultures of AA4 and M145 (Fig. 1E).

To determine the inducer and the producer, we cultured the strains independently in transwells, collected the conditioned medium from each isolated culture, and used that medium to grow the other strain. Addition of M145-conditioned medium induced robust antibiotic production by AA4 (Fig. 1G), but addition of AA4-conditioned medium to M145 did not induce antibiotic production (Fig. 1F). These results suggested that a M145-derived compound(s) stimulates the production of an antimicrobial agent by AA4.

Since antibiotic biosynthesis in *Actinomycetes* is intricately linked to carbohydrate metabolism we measured the carbohydrate content of M145-conditioned medium, AA4-conditioned medium, and fresh medium (22). Glucose was the predominant sugar in fresh medium (Fig. S2A). Conditioning of the medium by M145 reduced the amount of glucose and increased the concentration of galactose (Fig. S2A). Adding galactose, as well as *N*-acetyl-glucosamine (Glc*N*Ac), glucosamine, xylose, or arabinose, robustly induced antibiotic production in AA4 in the absence of M145 (Fig. S2B). This agrees with previous studies that found that Glc*N*Ac and xylose induce antibiotic synthesis in *Actinomycetes* through gene regulation (23, 24). Possible sources of galactose in the M145-conditioned medium are teichulosonic acid and polydiglycosylphosphate polysaccharide components of M145, accounting for up to one third of its cell wall dry mass (25, 26). Importantly, different *Actinomycete* species contain distinct cell-associated saccharide mixtures, a feature that has been used to classify them (27).

### Purification and absolute structure determination of amycomicin (AMY)

Bioassay-guided isolation of the crude extract from large-scale fermentation of AA4 on solid agar media containing Glc*N*Ac led to the purification of AMY. AMY has the molecular formula C_19_H_29_NO_5_, based on HR-q-ToFMS. Both the ^1^H and ^13^C NMR spectra suggested that AMY is a highly functionalized fatty acid with carbon chemical shifts indicating a carboxylic acid (*δ*_C_ 174.7), a ketone (*δ*_C_ 210.5), an olefin (*δ*_C_ 123.7 and 132.7), and three oxygenated sp^3^ carbons (δ_C_ 65.9, 67.2, and 67.6).

Analysis of AMY’s 2D NMR spectra (HSQC, H2BC, COSY, HMBC, and ROESY) revealed a relationship to two previously reported metabolites, YM-47515 and aerocyanidin (28, 29). Both contain a highly unstable epoxide isonitrile functionality at the end of the fatty acid chain. Further analysis of AMY’s ^13^C NMR spectra revealed a small peak at 162.0 ppm (isonitrile carbon) that had no correlations in any of the 2D NMR spectra. Additionally, infrared spectra on AMY consisted of peaks at 3350, 2133, and 1704/1651 cm^-1^, which indicate a hydroxyl/carboxylic acid, a triple bond, and two carbonyl groups, respectively. Thus, analysis of the NMR dataset and comparison to both YM-47515 and aerocyanidin led to the planar structural assignment of AMY as shown in Fig. 2, a highly modified fatty acid containing an epoxy-isonitrile warhead. Amycomicin inhibits growth of *Staphylococcaceae* at 30 nM and does not inhibit other tested microbes (Fig. S9)

**Fig. 2.**
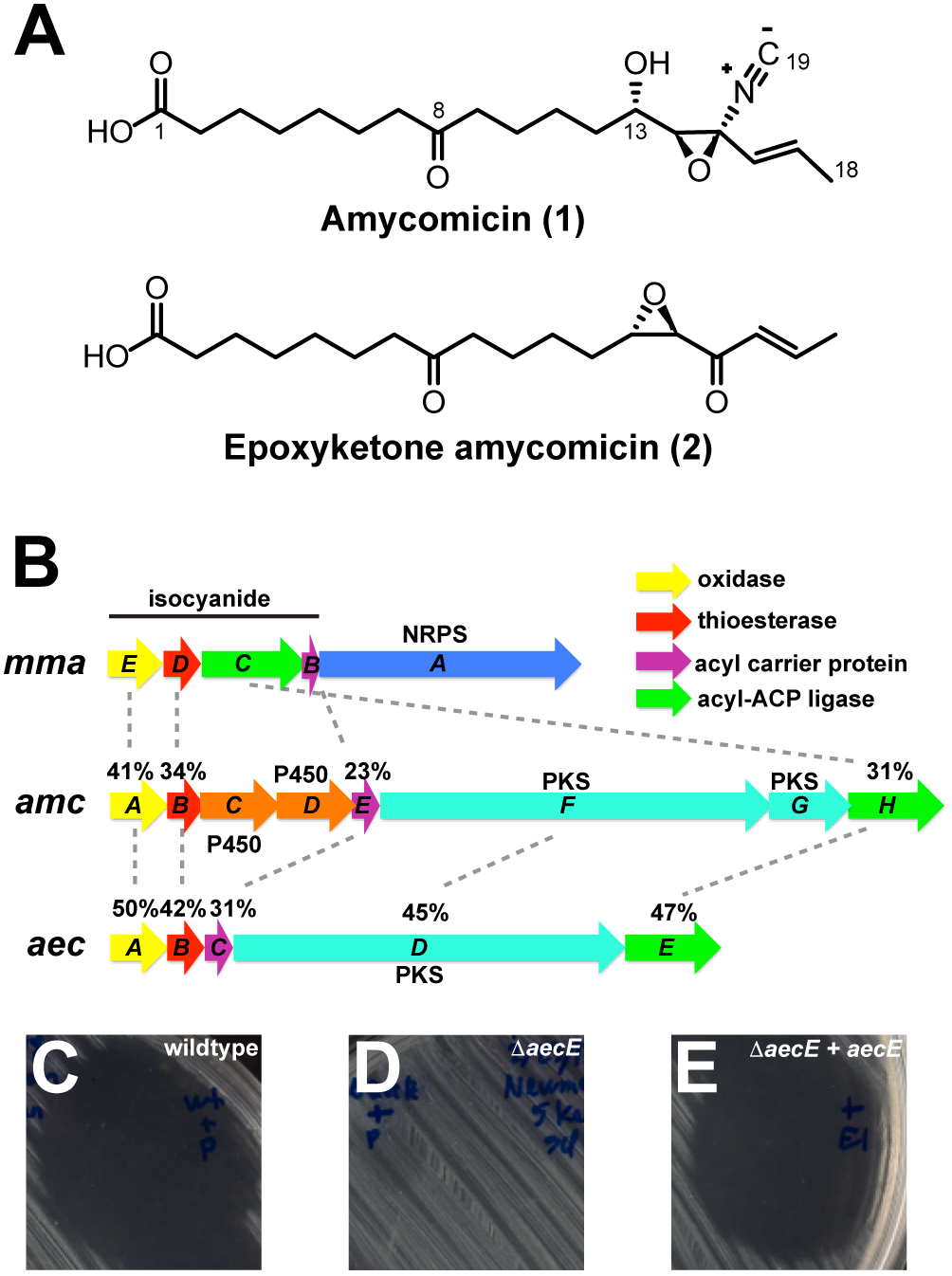
Molecular structure and biosynthetic gene cluster of amycomicin. (*A*) Molecular structures of amycomicin (1) and epoxyketone amycomicin (2). (*B*) Gene conservation between BGCs of isonitrile lipopetide from *Mycobacterium marinum* (*mma*), amycomicin from AA4 (*amc*), and aerocyanidin from *Chromobacterium* sp. ATCC 53434 (*aec*). Percentages indicate amino acid sequence identity. (*C-E*) Halo (or lack thereof) produced within a lawn of *S. aureus* by 10 μL of filtered supernatant from wildtype ATCC 53434 (*C*) or Δ*aecE* (*D*) containing an empty expression plasmid, and Δ*aecE* containing a plasmid expressing *aecE* (*E*).

Epoxy-isonitrile functional group is quite labile, therefore its decomposition in basic media was used as part of our structural analysis and it likely forms a key feature in AMY’s mechanism of action. Relative stereochemistry of AMY was assigned to be *syn* by a key ROESY correlation between H14 and H16. The relative C13 hydroxyl stereochemistry was determined by converting AMY to epoxy-AMY with inversion at C14 (30), which was subsequently assigned as a *trans*-epoxide because of the measured 2 *Hz* coupling between H13 and H14. Conversion of AMY to epoxy-AMY releases a molecule of HCN and leaves behind an epoxy ketone, which is a familiar component of naturally occurring biologically active molecules (31). The absolute stereochemical assignment of epoxy-AMY was assigned by comparing its CD Cotton effect and specific optical rotation to known epoxy-ketone containing metabolites (32-34).

### Identification of a putative amycomicin biosynthetic gene cluster

The isonitrile moiety led to the identification of AMY’s putative biosynthetic gene cluster. The isonitrile function is uncommon in natural products, but two different genes have been identified in bacteria that are able to install this moiety. The first isonitrile synthase (IsnA) to be identified utilizes ribose as a source of the isonitrile carbon, which is attached to an existing amine (35, 36). AA4 has no IsnA homologs. In the second pathway, an isonitrile moiety is created on an existing acyl chain via the action of four proteins, which were recently discovered to install the isonitrile functionality in a lipopeptide produced by *Mycobacterium marinum* (19). We found homologs of *mmaB-E* clustered within AA4 genome (Fig. 2B). We named this gene cluster *amc*. In addition to genes predicted to synthesize isonitrile (*amcA, B, E* and *H*), this cluster also contains genes encoding putative polyketide synthases (*amcF, G*) and two cytochrome P450s (*amcC, D*), which could be responsible for the biosynthesis of the AMY’s polyketide backbone and epoxide, respectively.

As we were unable to generate mutations in the genome of AA4, we sequenced the genome of *Chromobacterium* sp. ATCC 53434, which produces aerocyanidin, another member of the epoxide isonitrile antibiotic family (29). In the genome of ATCC53434 we found a gene cluster (which we named *aec*) that is very similar to *amc* (Fig. 2B). A mutant of ATCC 53434’s putative acyl-ACP-ligase gene (Δ*aecE*) abolished production of aerocyanidin and its supernatant did not inhibit *S. aureus*. Production of aerocyanidin was restored by supplementation of *aecE in trans* (Fig. 2C-E, and S5). Given the clear structural similarities between aerocyanidin and AMY, linking of aerocyanidin biosynthesis to the cluster provides strong evidence that the similar gene cluster within AA4 genome is responsible for biosynthesis of AMY.

### Fatty acids protect *S. aureus* against amycomicin

Serendipitously, we found that another AA4 metabolite protects *S. aureus* from AMY. We sometimes observed that AA4-conditioned medium led to a faint zone of growth <u>within</u> the zone of inhibition produced by AMY (Fig. 3A), which suggested that AA4 could produce a factor that served as an ‘antidote’ to AMY. We purified this factor from AA4-conditioned medium using high performance liquid chromatography (HPLC) tracking anti-AMY activity. This strategy identified a compound that allowed *S. aureus* growth in AMY’s presence (compare Fig. 3B and 3C). Mass spectrometry (MS), NMR analysis, and gas chromatography MS identified the compound as palmitoleic acid, a 16-carbon straight-chain unsaturated fatty acid (SCUFA – Fig. S6). Commercially available palmitoleic acid as well oleic acid, an 18-carbon fatty acid allowed growth of *S. aureus* in the presence of AMY (Fig. 3D and 3E).

**Fig. 3.**
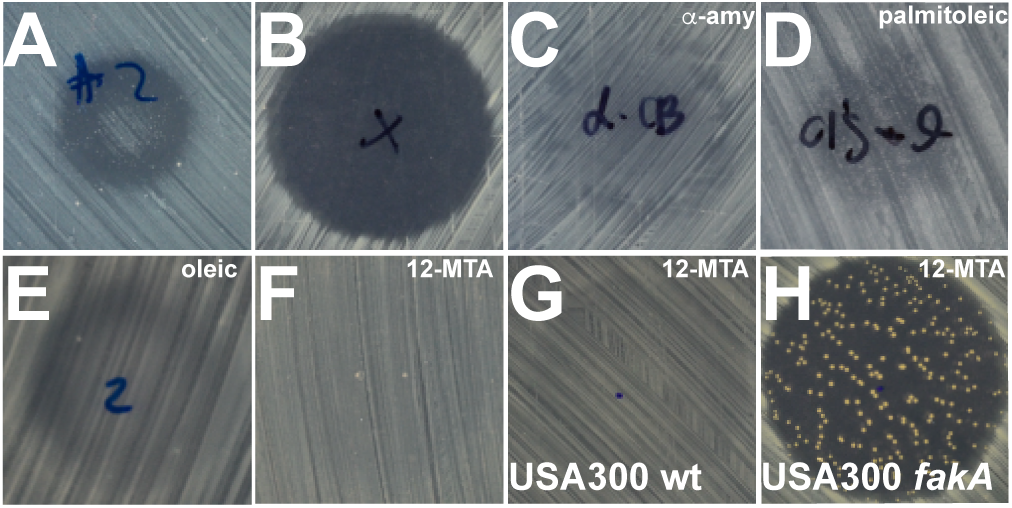
Reversal of amycomicin activity by fatty acids. (*A*) Small zone of growth within a zone of growth inhibition produced by a batch of AA4-conditioned medium. (*B-F*) Halos (or lack thereof) produced by (*B*) 100 ng amycomicin, (*C*) amycomicin + “anti-amycomicin”, (*D*) amycomicin + palmitoleic acid, (*E*) amycomicin + oleic acid, or (*F*) amycomicin + 12-MTA. (*G* and *H*). Results of adding amycomicin + 12-MTA to a lawn of wild-type *S. aureus* USA300 (G) and to a lawn of USA300 *fakA* transposon mutant (H).

S. *aureus* does not synthesize unsaturated fatty acids such as oleic acid or palmitoleic acid, but it can incorporate them from the medium (37, 38). Instead, *S. aureus* synthesizes branched chain fatty acids (BCFAs) that function similarly to SCUFAs (39). The most abundant BCFA produced by *S. aureus* is 12-methyltetradecanoic acid (12-MTA, anteiso-15:0). We found that addition of 12-MTA also inhibits AMY’s activity (Fig. 3F). Addition of increasing amounts of either 12-MTA or oleic acid protected *S. aureus* from AMY in a dose-dependent manner up to a concentration where the fatty acids become toxic (Fig. S7A and S7B). In contrast, straight-chain saturated fatty acids (SCSFA) do not protect *S. aureus* against amycomicin (Fig. S7C and S7D).

Incorporation of external fatty acids into the membrane of *S. aureus* requires fatty acid kinase A (FakA) (37). A *S. aureus* mutant with a transposon insertion in the gene encoding FakA (*fakA*) was sensitive to AMY even in the presence of 12-MTA (Fig. 3G and 3H). We noticed frequent emergence of AMY-resistant mutants in the *fakA* mutant background when in the presence of 12-MTA. These mutants reverted to a wild-type phenotype where they gained the ability to utilize 12-MTA to counteract AMY, but were still susceptible to AMY in the absence of added fatty acids (Figs. S6E and S6F). These observations pointed to the possibility that AMY targets fatty acid biosynthesis. Interestingly, a number of fatty acid biosynthesis genes are located adjacent to *amc* (Fig. S11).

### Amycomicin alters cell morphology of *S. aureus*

To investigate the effect of AMY on cell morphology we utilized transmission electron microscopy on ultrathin-sectioned *S. aureus*. Compared to untreated cells, bacteria treated with AMY displayed gross alterations. Specifically, the cells contained thickened cell walls and a severely shrunken cytoplasmic compartment (compare Fig. 4A and 4B). Similar phenotypes were observed in cells treated with the fatty acid biosynthesis inhibitors platensimycin and irgasan (Fig. 4C and 4D, respectively) (40, 41). These results indicate that AMY alters cell morphology in a manner similar to known fatty acid biosynthesis inhibitors.

**Fig. 4.**
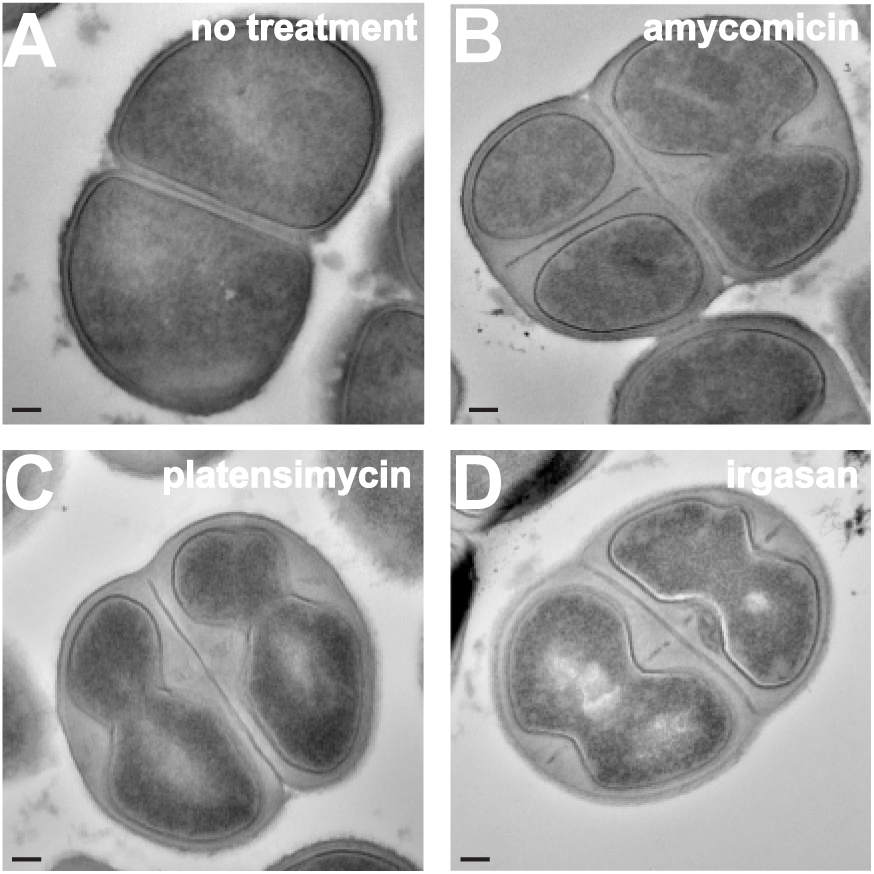
Morphological changes in *S. aureus* induced by treatment with amycomicin. Transmission electron micrographs of sectioned *S. aureus* cells that were (*A*) not treated with antibiotics or treated with (*B*) amycomicin, (*C*) platensimycin, or (*D*) irgasan. Scale bar is 100 nm.

### FabH overexpression increases resistance to AMY

It seemed likely that AMY targets an enzyme involved in fatty acid biosynthesis, therefore we manipulated expression levels of fatty acid biosynthetic enzymes, an approach that has previously been used to identify antibiotic targets (42-45). As a control, we cloned the 3-oxoacyl-[acyl-carrier-protein] synthase 2 gene (*fabF*) into a plasmid under a constitutive promoter and tested its effect on susceptibility of *S. aureus* to platensimycin, a well-characterized FabF inhibitor (40), and observed a 4-fold increase in resistance to platensimycin (Fig. S8). AMY has a minimal inhibitory concentration of ~30 nM (~10 ng/mL) against *S. aureus* grown at 37°C at pH 7.0. In assays against strains overexpressing individual Fab proteins, overexpression of 3-oxoacyl-[acyl-carrier-protein] synthase 3 (FabH) displayed a 4-fold increase in the MIC against AMY while overexpression of other Fab proteins had no effect (Fig. 5A).

**Fig. 5.**
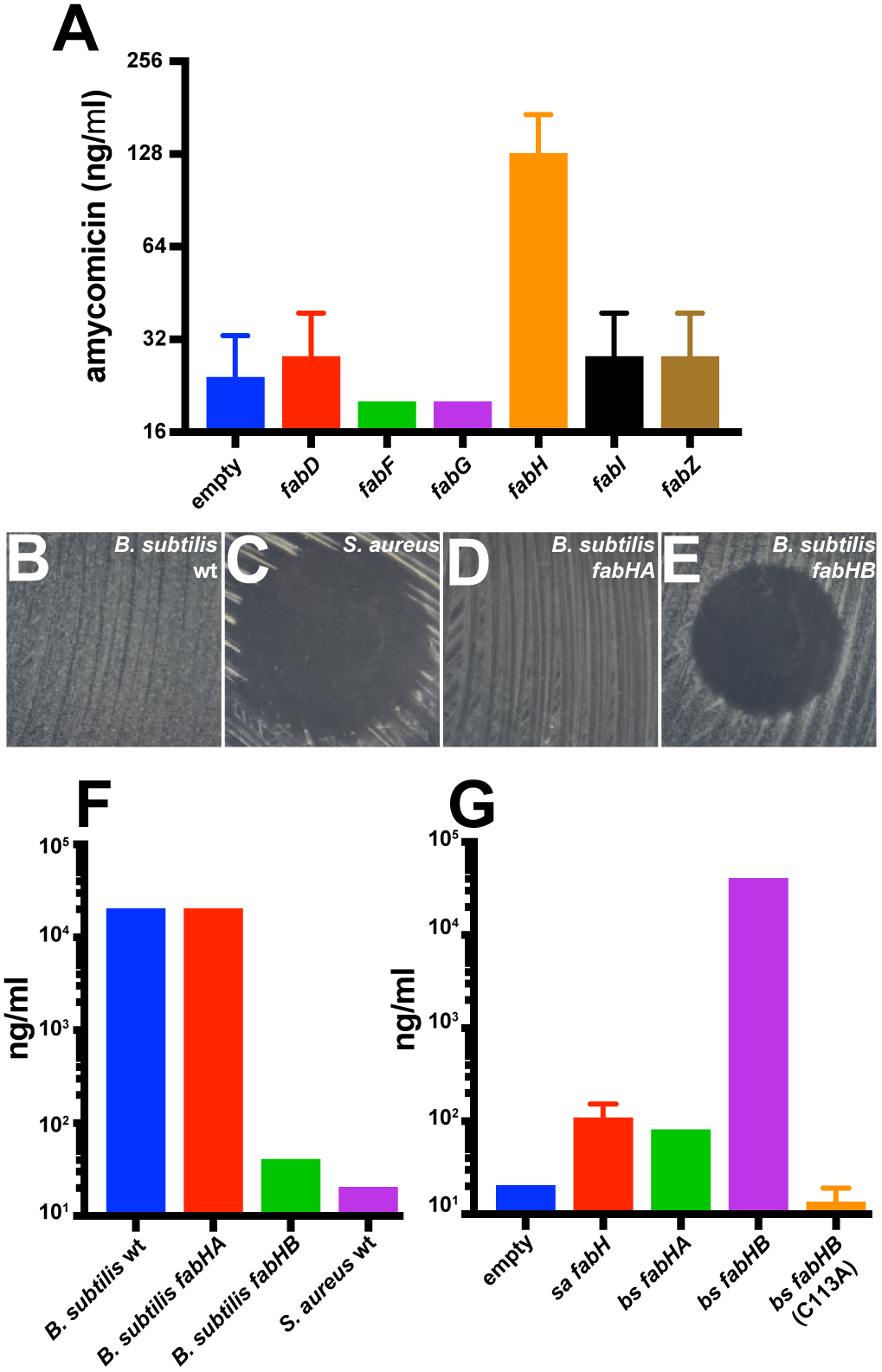
Protection against amycomicin inhibition in *S. aureus* by FabH overexpression. (*A*) Minimal inhibitory concentrations of amycomicin on strains of *S. aureus* carrying a plasmid overexpressing individual *S. aureus* fatty acid biosynthesis genes. (*B-E*) Effect of adding 100 ng amycomicin to lawns of (*B*) wild-type *B. subtilis,* (*C*) wild-type *S. aureus,* (*D*) *B. subtilis fabHA,* and (*E*) *B. subtilis fabHB* (*F*). Minimal inhibitory concentrations of amycomicin against the indicated strains of *B. subtilis* and *S. aureus* (*G*). Minimal inhibitory concentrations of amycomicin *S. aureus* strains harboring the indicated plasmids, that led to the overexpression of the native *S. aureus* FabH or the *B. subtilis* FabHA or FabHB. Error bars represent standard deviations.

### *Bacillus subtilis* FabHB confers resistance against AMY

Members of the *Staphylococcacceae* family are uniquely susceptible to AMY (Fig. S9). We employed *Bacillus subtilis* that is resistant to AMY (compare Fig. 5B and 5C) in order to understand this unique susceptibility. *B. subtilis* encodes two homologs of FabH and thus single-knockout mutants are viable (46, 47). FabHA is more closely related to *S. aureus* FabH (58% identity, 76% similarity) than FabHB (42% identity, 59% similarity) (47). *B. subtilis fabHA* is resistant to amycomicin, but *B. subtilis fabHB* is sensitive (compare Fig. 5D and 5E). The MIC measurements indicated a nearly three orders of magnitude decrease in resistance to AMY of *B. subtilis fabHB* knockout compared to wild-type *B. subtilis* or *B. subtilis fabHA* knockout (Fig. 5F). Further, expressing *fabHB* from *B. subtilis* in *S. aureus* resulted in a 500-fold increase in the MIC of AMY, whereas expression of *fabHA* did not (Fig. 5G). Expression of a catalytically inactive C113A mutant of FabHB did not increase resistance of *S. aureus* to AMY. Collectively these results identify *S. aureus* FabH as AMY’s target. It is important to note in regard to our original goal of developing a targeted interaction screen, that a previous high-throughput screen identified all natural product inhibitors of FabH known to date, but did not discover AMY (48).

### AMY alters fatty acid composition of *S. aureus*

FabH’s preference for branched-chain or straight chain acyl-coenzyme A substrates determines the membrane BCFA/SCSFA ratio (46). *S. aureus* growing in exponential phase were treated with a range of concentrations of AMY. In untreated cells, 12-MTA (anteiso-15:0) was the most abundant fatty acid (Table S3). *S. aureus* cells treated with AMY displayed an altered BCFA/SCFA ratio with a decrease of a total fraction of all BCFAs (from 77% to 46%) and 12-MTA specifically (from 37% to 22%, Figure S7G and Table S3). Consistent with this observation thiolactomycin, a known inhibitor of FabH, also reduces BCFA content in *Streptomyces collinus* (49). These results are also consistent with the observation that addition of BCFAs reverses the activity of AMY (Fig. 3).

### AMY protects against *S. aureus* infection in a mouse skin model of infection

To explore the potential of AMY as a therapeutic agent we used a superficial mouse skin infection model (50). *S. aureus* USA300 containing a modified *lux* operon allowed us to monitor the progression of the infection in live animals (51). The infection developed untreated for 24 hours, and the mice were separated into three groups based on the treatment to be received. Bioluminescence was measured to ensure that the mice in the three groups had similar burdens of infection (Fig. S10A). Mice in the first group were left untreated (negative control), mice in the second group were treated with a topical formulation of 2% mupirocin, and mice in the third group were treated with 0.001% solution of AMY once a day for three days. The mice treated with AMY for three days showed a marked decrease in luminescence compared to untreated mice (Fig. 6A and 6B). The decrease was similar to that observed in infection sites treated with mupirocin. The decrease was confirmed by the reduction in number of colony forming units isolated from the infection sites (Fig. S10B). These data demonstrate that AMY has *in vivo* activity in a mouse model of *S. aureus* skin infections. While AMY’s potency and specificity against *S. aureus* are highly desirable, it’s lability makes it an unlikely candidate to advance along a therapeutic pipeline. Although AMY resulted in decreased infection, in our initial study it acted slower than mupirocin (Fig. 6A, 6B, S10C, S10D). While mupirocin treatment resulted in a significant decrease in infection one day after treatment, amycomicin did not have an effect until day two. However, mupirocin used as a control in this experiment was applied in the form of a pharmaceutical-grade ointment at a 2% concentration, a concentration much higher than the amycomicin concentration (0.001%) we used.

**Fig. 6.**
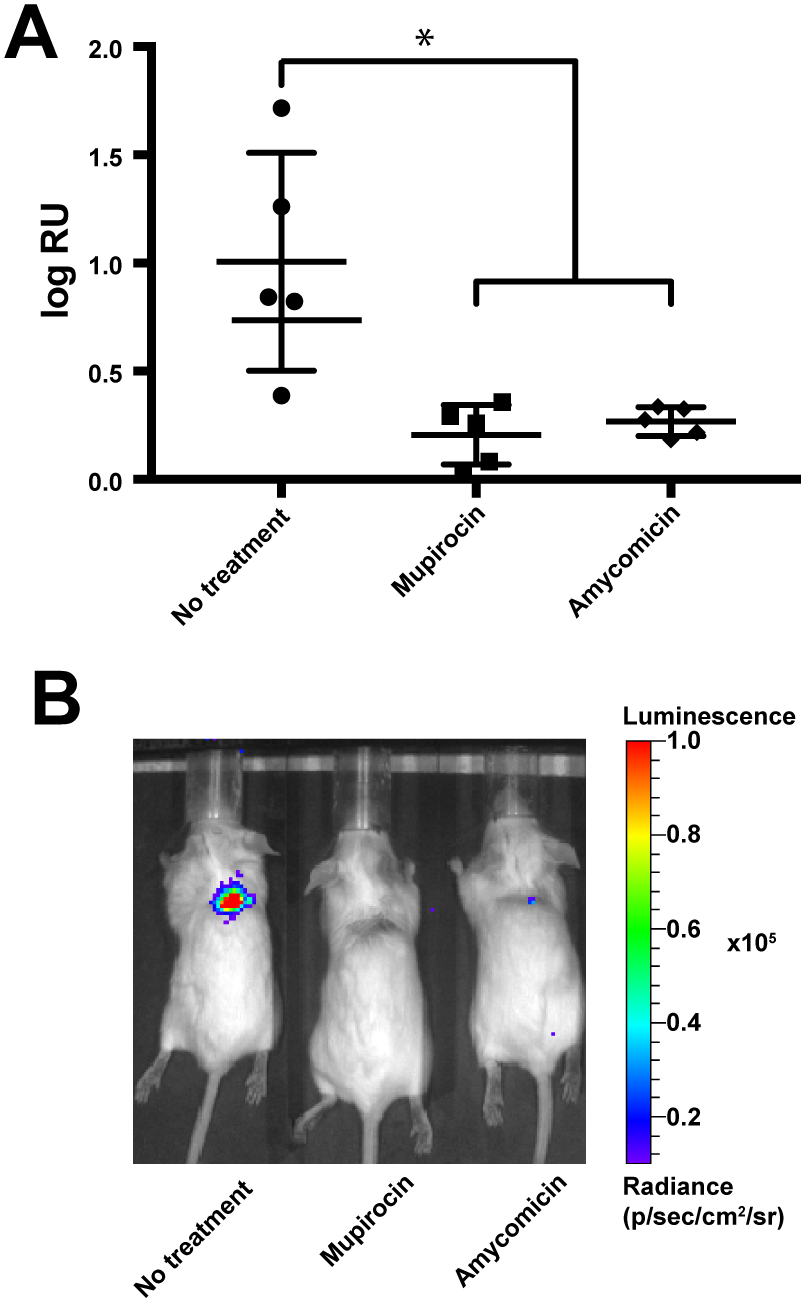
Murine *S. aureus* skin infections and treatment. (*A*) Bioluminescence expressed as radiance units (RU) detected in the infected sites 4 days post-infection and after 3 daily doses of the indicated treatment. Horizontal bars indicate averages and standard deviations of log_10_-transformed values. Welch’s t-test was used to determine the p-values. P-values that are smaller than 0.05 are marked with an asterisk. (*B*) Representative images of bioluminescence emanating from infected sites after three doses of the indicated treatment.

### Conclusion

In an effort to discover new antibiotics we designed and tested an approach coupling the ability of bacterial co-cultures to elicit the expression of cryptic metabolites with phenotypic screening against an important human pathogen. The screen exploits the stochastic populations produced by identical but complex starting sets, and the ability of rapid 16S rDNA sequencing to deconvolute and identify productive populations. While this initial targeted interaction approach to discovering new antibiotics was successful, it is important to note a paradox that was troubling from the outset: the logic of the interaction screen is based on taking advantage of existing environmental signals. AA4 is a soil *Actinomycete* and as such is not likely to encounter *S. aureus*, a common inhabitant of humans. AMY does not inhibit M145, *B. subtilis*, or any other soil microbe we tested it against. While we do not understand the ecological role of AMY, our results are encouraging for future applications of target-based interaction screening as the starting community need not reflect inhabitants of the target’s community. Similar screens can theoretically be applied to any cultivable complex multispecies community, such as host-associated microbiota.

## Acknowledgments

We thank members of the R.K. and J.C. laboratories for valuable advice. This work was supported by NIH grants GM058213, R21AI117025 (RK), and R01AT009874 (JC). We thank the Harvard Medical School Center for Macromolecular Interactions, ICCB-Longwood Screening Facility, Small Molecule Mass Spectrometry Core Facility at Harvard Univeristy, and East Quad NMR Facility for analytical services. SK was supported by the Intramural Research Program of the National Human Genome Research Institute, National Institutes of Health. This work utilized the computational resources of the NIH HPC Biowulf cluster (https://hpc.nih.gov). Carbohydrate analysis was supported by the Chemical Sciences, Geosciences and Biosciences Division, Office of Basic Energy Sciences, U.S. Department of Energy grant (DE-FG02-93ER20097) to Parastoo Azadi at the Complex Carbohydrate Research Center.

## SI Methods

### General experimental procedures

Specific optical rotation data was measured on a JASCO P-2000 polarimeter, circular dichroism spectra on a JASCO J-815, UV spectra on an Amersham Biosciences Ultrospec 5300-pro UV/Visible spectrophotometer, and IR spectra on a Bruker Alpha-p. NMR data were acquired on a ~2 mg sample of **1** or less than 0.5 mg sample of **2** dissolved in 35 μL *d*_6_-DMSO or CD_3_OD, respectively, using either 500 MHz Varian VNMRS (Varian NMR System) equipped with an inverse cold probe, Bruker 500 MHZ spectrometer equipped with a broadband cyroprobe or Bruker AVANCE II spectrometer equipped with a ^1^H{^13^C/^15^N} cryoprobe. LR-LCMS data were obtained using an Agilent 1200 series HPLC system equipped with a photo-diode array detector and a 6130-quadrupole mass spectrometer. HR-ESIMS was carried out using an Agilent LC-q-TOF Mass Spectrometer 6530-equipped with a 1290 uHPLC system. HPLC purifications were carried out using Agilent 1100 or 1200 series HPLC systems equipped with a photo-diode array detector. All solvents were of HPLC quality.

### Growth media

Due to the pH-dependence of amycomicin activity all experiments were carried out in buffered 1X PTSYM medium unless noted otherwise (1X PTSYM = 1 g/L Bacto^TM^ Peptone, 1 g/L BactoTM Tryptone, 1 g/L Bacto^TM^ Soytone, 1 g/L Bacto^TM^ Yeast Extract, 1 g/L Bacto^TM^ Malt Extract, 100 mM MOPS pH7.0). Solid media contained 1.5% agar. Susceptibility tests were carried out with *S. aureus* Newman at 37°C unless noted otherwise.

### Discovery of amycomicin using a transwell system

*Amycolatopsis sp.* AA4, *Streptomyces coelicolor* M145, *Streptomyces sviceus* ATCC 29083, *Kutzneria sp.* 744, *Micromonospora sp.* M42, *Streptomyces sp.* Mg1, *Streptomyces sp.* E14, *Streptomyces hygroscopicus* ATCC 53653*, Streptomyces pristinaespiralis* ATCC 25486 were used to construct the *Actinomycete* community. Approximately 10^6^ spores of each species were inoculated onto transwells (Millipore, Billerica, MA; PIHT24H48) containing 100 *μ*l 0.75% agarose (MP Biomedicals, Santa Ana, CA 820721) either in isolation or as a mix totaling approximately 10^7^ spores. The bottom well (24-well plate) of each sample contained 1 ml of 0.1X PTSYM medium composed of 0.01% peptone, 0.01% tryptone, 0.01% soytone, 0.01% yeast extract and 0.01% malt extract broth and incubated at 30 °C. Stacks of multi-well plates containing the transwells were placed inside plastic containers to slow down evaporation. The broth from the bottom wells was tested every 1-2 days for antibiotic activity by spotting 10 *μ*l on agar plates with *S. aureus* Newman. After 3 weeks, the transwells were transferred onto wells containing fresh medium. The antibiotic activity became detectable a day later in a nine-member community. For a bipartite community AA4 and M145 were both inoculated at approximately 10^6^ spores each and the experiment was carried out as for a multispecies community. For the remainder of the experiments the transwell size was increased to a 6-well plate format to increase yields (Greiner Bio-One, Kremsmünster, Austria; 657640). These transwells contained 2 ml of agarose and the bottom wells contained 4 ml medium. The medium also contained 10 mM MOPS pH7.0 to control the pH and improve reproducibility. For AA4-M145 cross-inductions, the two strains were grown separately. After 3 weeks, the transwells were transferred onto fresh media, which was conditioned for another 1-2 days. The transwells containing AA4 where then placed into media preconditioned by M145, and transwells containing M145 where placed into media preconditioned by AA4. The antibiotic activity was observed 1-2 days later in media that was conditioned sequentially by M145 and AA4, but not *vice versa*.

### Sequencing of 16S rDNA gene

Two hundred microliter agarose plugs containing *Actinomycete* communities were stored at −80 °C until further processing. The plugs were placed inside 2 ml vials and combined with 500 *μ*l of 0.1 mm zirconia/silica beads (BioSpec Products, Bartsville, OK), 500 *μ*l buffer ATL containing 5 *μ*l RNAse A and 500 *μ*l buffer AL (Qiagen, Hilden, Germany). The samples were homogenized twice for 45 seconds using FastPrep FP120 (Thermo Scientific Savant, Waltham, MA). The beads and cell debris were pelleted for 15 minutes at 15,000 × *g* and 800 *μ*l the supernatant was mixed with 40 *μ*l proteinase K (Qiagen, Hilden, Germany) and incubated at 70°C for 30 minutes. Forty microliters of 3M sodium acetate (pH5.2) were added to the samples and the pH of the sample confirmed to be < 7.0. Afterwards, the samples were mixed with 440 *μ*l ethanol. The samples were loaded onto columns from Qiagen DNeasy Blood & Tissue Kit and purified further following the kit instructions. DNA concentration was measured by Quant-IT^™^ dsDNA Assay Kit and diluted to 1 ng/ml. The V4 region of the 16S rDNA gene was amplified over 25 cycles and sequenced as previously described using MiSeq Reagent Kit v. 2 500 cycles (Illumina, San Diego, CA) (1). The data were analyzed with QIIME (MacQIIME 1.8.0) pipeline software (1). UCHIME was used to remove chimeric sequences (2). Due to high similarity between the 16S rDNA sequences of the community members, the reads were binned based on 99% identity using uclust, using a custom reference library containing the 16S rDNA sequences of the community members (3).

### Glycosyl composition analysis of the media

Ten milliliters of fresh medium or media conditioned by M145 or AA4 as described above was lyophylized and analyzed for glycosyl composition at the Complex Carbohydrate Research Center at the University of Georgia. Glycosyl composition analysis was performed by combined gas chromatography/mass spectrometry (GC/MS) of the per-*O*-trimethylsilyl (TMS) derivatives of the monosaccharide methyl glycosides produced from the sample by acidic methanolysis (4). Briefly, the samples (300ug) were heated with methanolic HCl in a sealed screw-top glass test tube for 17 h at 80 °C. After cooling and removal of the solvent under a stream of nitrogen, the samples were treated with a mixture of methanol, pyridine, and acetic anhydride for 30 min. The solvents were evaporated, and the samples were derivatized with Tri-Sil^®^ (Pierce) at 80 °C for 30 min. GC/MS analysis of the TMS methyl glycosides was performed on an Agilent 7890A GC interfaced to a 5975C MSD, using an Supelco Equity-1 fused silica capillary column (30 m × 0.25 mm ID).

### Induction of amycomicin by individual sugars

Transwells inoculated with AA4 were placed inside bottom wells containing defined medium (2 mM MgCl_2_, 0.15 mM CaCl_2_, 6.7 mM KCl, 10 mM NH_4_Cl, 10 *μ*M FeCl_3_, 3 mM pyruvate, 1/100 dilution of ATCC Vitamin Supplement, 10 mM MOPS pH7.0). After three weeks, the old medium was replaced with the same fresh medium supplemented with 1 mM sugars. Three days post-induction 10 *μ*l of AA4-conditioned media were two-fold serially diluted and spotted on standard 90 mm Petri dishes containing 15 ml 1X PTSYM (buffered with 100 mM MES, pH6.0). Once the spots were absorbed, *S. aureus* was spread on these plates with cotton-tipped applicators and incubated at 22 °C. The amount of relative activity was assessed in arbitrary units quantified by determining the highest dilution factor that allowed for visible inhibition of *S. aureus* Newman.

### Amycomicin purification

Spores of AA4 were spread with cotton applicators onto media containing 1.5% agar, 50 mM *N*-acetyl-glucosamine (Sigma, St. Louis, MO) and 10 mM MOPS pH7.0 totaling approximately 8 liters. The Petri dishes inoculated with AA4 were incubated at room temperature for 2-4 weeks in plastic containers. Agar was then extracted twice with ethyl acetate for 4 hours at room temperature. Ethyl acetate was removed by evaporation under vacuum at room temperature to yield a crude extract. In order to prevent degradation of the active metabolite, the crude extract was stored at −20 °C under an inert atmosphere (argon). The crude extract was fractionated using reverse-phase C18 Sep-Pak columns (Waters, Milford, MA) and eluted with a stepwise gradient (0%, 25%, 50%, 75% and 100% acetonitrile/water) resulting in five fractions. Bioassay on the four SPE fractions showed the activity was predominately in the middle polarity fractions – 50% and 75% acetonitrile/water fractions. The active fractions were dried down under vacuum with gentle heating and subsequently resuspended in 50% acetonitrile/water to 10 mg/ml and filtered for further purification by high performance liquid chromatography. Further fractionation using a Phenomenex 4 μm Hydro semi-preparative column, holding 40% ACN + 0.1% formic acid (FA)/60% H_2_O + 0.1% FA for 5 min then a gradient to 55% ACN + 0.1% FA/45% H_2_O + 0.1% FA over 30 min, yielding pure amycomicin (Fig. 2A).

Amycomicin (**1**): colorless oil; [α]^22^ −18.7 (*c* 0.43, MeOH); UV (MeOH) *λ*_max_ (log ε) 209.0 (3.71), 302.0 (2.90) nm; IR (neat) *v_max_* 3350, 2932, 2859, 2133, 1704, 1651, 1454 cm^-1^; NMR (500 MHz, CD_3_OD) and ^13^C NMR (125 MHz, CD_3_OD) see table S1; HRESIMS [M+H]^+^ 352.2119 *m*/*z* (calcd for C_19_H_30_NO_5_ 352.2119, Δ 0.0 ppm).

Epoxyketone amycomicin (Fig. 2B): pale yellow oil; [α]^22^ −13.3, (*c* 0.32 MeOH); UV (MeOH) *λ*_max_ (log ε) 229.0 (2.71) nm; IR (neat) *v_max_* 2930, 2850, 1698, 1633, 1416 cm^-1^; NMR (600 MHz, CD_3_OD) and ^13^C NMR (150 MHz, CD_3_OD) see table S2; HRESIMS [M+H]^+^ 325.2013 *m*/*z* (calcd for C_18_H_29_O_5_ 325.2015, Δ 0.6 ppm).

### Stereochemical analysis of amycomicin

An aliquot of purified amycomicin (~1 mg) was resuspended in 0.5 mL of 0.05 M sodium phosphate buffer, pH 11.4, for 5 min at room temperature (5). The pH of the reaction mixture was adjusted to approx. 4.5 with 80 μL of a 1 M phosphoric acid solution (aq) and then this mixture was extracted with ethyl acetate (1 mL x 3). The combined organic layer was subsequently washed with brine (1 mL), dried with MgSO_4,_ and concentrated under vacuum. Analytical data, including HRMS and 2D NMR dataset, confirmed that the predominate product was compound **2**, where HCN was eliminated. Both the circular dichroism (CD) spectrum and specific optical rotation of **2** revealed a negative cotton effect. Comparison to literature values led to the absolute assignment of **2** as 13*S*,14*R* and thus 13*S*,14*S*,15*S* stereochemistry for amycomicin (**1**).

### Genome sequencing and assembly

*Amycolatopsis sp.* AA4 genome was sequenced with PacBio single-molecule real-time (SMRT) sequencing technology at the University of Delaware DNA Sequencing and Genotyping Center. Five micrograms of genomic DNA were sheared using g-tube (Covaris) to 20kb fragments. The Pacbio library was prepared using standard Pacbio protocol for 20 kb libraries “20 kb Template Preparation Using BluePippin Size-selection system”. After Blue Pippin (Sage Science) size selection from 10 kb average DNA fragments size of the library was 25 kb. Sample library was sequenced on Pacbio RS II instrument with one SMRT cell using P6-C4 chemistry with 6-hour movie. AA4 genome (accession numbers CP024894-CP024896) was assembled with Canu v1.4 +183 changes (r8178 82bf1d3e8b6e74f534fd7b80b24e84e66807a7bd) ran with default parameters (6). ATCC 53434 genome (accession number CP025429) was assembled with Canu v1.6. Self-alignments were computed with MUMmer3.23 and redundant sequence trimmed off the ends of the circular sequences (7). Arrow v2.1.0 was used to generate the final polished assembly (8). Protein IDs relevant to this publication are ATY12794.1-ATY12796.1 (AmcCBA), ATY16518.1 (AmcD), ATY16517.1 (AmcE), ATY12791.1-ATY12793.1 (AmcHGF); AUH51997.1 (AecA), AUH53650 (AecB), AUH51994.1-AUH51996.1 (AecEDC).

### *Chromobacterium* sp. ATCC 53434 mutagenesis and complementation

One kb regions flanking *aecE* were amplified using the primers listed in Table S4. PCR products were joined, inserted into pNPTS138 Cm plasmid (Acquired from Addgene, deposited by Howard Steinman Lab) and transformed using NEBuilder ^®^ HiFi DNA Assembly Cloning Kit. *E. coli* transformants were selected for chloramphenicol (20 *μ*g/ml) resistance, plasmid isolated, and transformed into *E. coli* S17-1 conjugation strain. For conjugation, S17-1 containing the plasmid was grown in LB supplemented with chloramphenicol while ATCC 53434 was grown in LB supplemented with ampicillin (100 *μ*g/ml) until early stationary phase. Cells were washed once in LB and resuspended in 1/10th volume of LB. One hundred *μ*l of S17-1 cells were mixed with 10 *μ*l of ATCC 53434, spread on LB agar and incubated at 30 °C overnight. The cells were then scraped off and spread on LB agar supplemented with chloramphenicol (to select for the plasmid) and ampicillin to which ATCC 53434 is naturally resistant. Because ATCC 53434 develops resistance to chloramphenicol several colonies were picked the next day and inoculated into LB broth containing ampicillin and incubated with shaking overnight at 30 °C. The overnight cultures were diluted 10,000 times and 100 *μ*l were spread on LB agar containing 100 *μ*g/ml ampicillin and 5% (w/v) sucrose. Resulting colonies were replica-plated on LB agar containing ampicillin with or without chloramphenicol. Isolates that lost chloramphenicol resistance were further screened for *ΔaecE* mutation and loss of the plasmid by PCR and sequencing.

To complement the mutation wild type *aecE* was PCR-amplified using the primers listed in Table S4 and inserted into plasmid pBBR1MCS-2 (9). *E. coli* transformants were selected for kanamycin (50 *μ*g/ml) resistance, plasmid isolated, transformed into *E. coli* S17-1, and conjugated into Δ*aecE*. Because this strain failed to produce aerocyanidin, we replaced *lac* promoter found on the plasmid with the native *aec* promoter (see Table S4). Empty pBBR1MCS-2 was similarly introduced into wildtype and Δ*aecE* to serve as controls.

### Aerocyanidin analysis

ATCC53434 was inoculated into 50 ml media containing 0.5% tryptone, 0.3% yeast extract, 0.3% malt extract, 1% *N*-acetyl-glucosamine and 100 mM MOPS pH7.0. Strains containing pBBR1MCS-2 construct were supplemented with 5 *μ*g/ml kanamycin. The cultures were incubated with shaking at 25 °C. After three-five days, the cultures were centrifuged at 10,000 *g* for 30 minutes and the supernatant was filtered. The filtrate was fractionated using reverse-phase C18 Sep-Pak columns (Waters, Milford, MA) and eluted with a stepwise gradient (0%, 25%, 75%, 100% acetonitrile/water) resulting in four fractions. An aliquot (5 *μ*L) of the 75% fraction was analyzed by HRLCMS for production of aerocyanidin using a Phenomenex Kinetex 2.6 *μ*m EVO C18 100 Å (100 × 2.1 mm) under the following LC method: hold 10% ACN + 0.1% FA/H_2_O + 0.1% FA for 1 min then gradient to 100% ACN + 0.1% FA over 9 min then hold 100% ACN + 0.1% FA for 3.5 min with a constant flow rate of 0.3 mL/min.

### Palmitoleic acid purification and identification

The crude extract was fractionated using reverse-phase Sep-Pak columns (Waters, Milford, MA) and eluted with a stepwise gradient (0%, 25%, 50%, 100% acetonitrile/water) resulting in four fractions. Bioassay on the four SPE fractions showed the activity was in the 100% acetonitrile/water fraction. The active fraction was dried down under vacuum with gentle heating and subsequently resuspended in 90% acetonitrile/water to 10 mg/ml and filtered for further purification by high performance liquid chromatography. Further fractionation using a Phenomenex 4 μm C18 semi-preparative column, holding 85% ACN + 0.1% formic acid (FA)/15% H_2_O + 0.1% FA for 5 min then a gradient to 95% ACN + 0.1% FA/5% H_2_O + 0.1% FA over 20 min, yielding pure product. For GC/MS analysis the extracted fatty acid and standards were suspended in methanol at 1 mg/ml, esterified with (trimethylsilyl)diazomethane for 15 minutes and dried down under vacuum. Dried down solids were solubilized in hexanes and submitted to Small Molecule Mass Spectrometry Facility at Harvard University. The GC/MS analysis was conducted with 1 μL of the sample injected and at the inlet helium carrier gas flow rate of 1 mL/min in the constant flow mode with a fused-silica capillary column of cross-linked DB-23 (30 m × 0.25 mm × 0.25 μm, Agilent, Santa Clara, CA) on Quattro micro GC Mass Spectrometer (Waters, Milford, MA) equipped with a Combi PAL autosampler (CTC Analytics, Zwingen, Switzerland) and a split/splitless injector. The inlet and transfer line temperatures were set at 250 °C, and the ion source temperature was set at 200 °C. The GC conditions were as follows: oven temperature program, 50 °C for 1 min, 13 °C/min to 205 °C, 3 °C/min to 210 °C, 50 °C/min to 250 °C; split ratio, 75:1. The electron impact (EI)-MS conditions were as follows: full scan m/z range, 29-450 Da at 2.1-18.39 min. All data were acquired and analyzed with Waters MassLynx V4.1 software package.

### Testing fatty acids for reversal of amycomicin activity

#### Plate assays

Fatty acids (Sigma-Aldrich, St. Louis, MO) were suspended in methanol to 1 mg/ml. After 10 *μ*l of amycomicin was absorbed by the agar, 10 *μ*l of a fatty acid solution was spotted onto the same area and was also allowed to absorb before *S. aureus* Newman was spread on agar as described previously. Fatty acid kinase A (*fakA*, SAUSA300_1119) transposon mutant in the USA300 LAC JE2 background was acquired from Nebraska Transposon Mutant Library through BEI Resources (strain NE229) and propagated in the presence of 5 *μ*g/ml erythromycin (10). Resistant colonies of *fakA* mutant that proliferated in the presence of amycomicin and 12-MTA were re-streaked on tryptic soy agar and confirmed to have retained erythromycin resistance.

#### Liquid cultures

Overnight liquid cultures were diluted 1,000-fold into fresh 1X PTSYM and incubated for 2.5 hours. The cultures were then dispensed at 100 *μ*l per well into 96-well plates, supplemented with indicated concentrations of fatty acids and amycomicin and incubated for another 5 hours. Colony-forming units per milliliter of the media (CFU/ml) were measured at the time of inoculation, at the time of amycomicin/fatty acid addition and at the end of incubation. CFU/ml was quantified by serial dilution and plating on solid media.

### Cellular fatty acid content determination

To determine the effect of amycomicin on fatty acid content, 1 ml of *S. aureus* overnight cultures were subcultured into 270 ml of fresh media and incubated with shaking at 37 °C until OD600 reached approximately 0.1. At that point, the cultures were separated into 50 ml samples and amycomicin was added at indicated concentrations. These cultures were incubated for additional 2 hours, at which point the bacterial cells were pelleted by centrifugation at 8,000 *g* and frozen at −80 °C. The samples were submitted on dry ice to Microbial ID, Inc. (Newark, DE) where they were processed as described previously (11). Samples were analyzed using Agilent/HP 6890 Series II GC.

### Transmission electron microscopy

Overnight cultures of *S. aureus* were diluted to OD600 = 0.01, incubated until the OD600 reached 0.1 and separated into 2 ml aliquots in 13 × 100 mm glass test tubes. Amycomicin, platensimycin (Cayman, Ann Arbor, MI) and irgasan (Sigma, St. Louis, MO) were added at 100 ng/ml, 2.5 *μ*g/ml, and 30 ng/ml respectively. After 2 hours of incubation a 2x fixative was added to cultures to a final concentration of 2.5% Glutaraldehyde 1.25% Paraformaldehyde and 0.03% picric acid in 0.1 M sodium cacodylate buffer (pH 7.4) and pelleted by centrifugation. The samples were further processed and visualized at Harvard Medical School Electron Microscopy Facility. The pellet was refrigerated overnight, washed in 0.1M cacodylate buffer and post-fixed with 1% Osmiumtetroxide (OsO4)/1.5% Potassiumferrocyanide (KFeCN6) for 1 hour, washed twice in water, once in Maleate buffer (MB) and incubated in 1% uranyl acetate in MB for 1 hour followed by 2 washes in water and subsequent dehydration in grades of alcohol (10 minutes each; 50%, 70%, 90%, 100% twice). The samples were then put in propyleneoxide for 1 hour and infiltrated overnight in a 1:1 mixture of propyleneoxide and TAAB Epon (Marivac Canada Inc. St. Laurent, Canada). The following day the samples were embedded in TAAB Epon and polymerized at 60 °C for 48 hours. Ultrathin sections (about 60 nm) were cut on a Reichert Ultracut-S microtome, picked up on to copper grids stained with lead citrate and examined in a JEOL 1200EX Transmission electron microscope or a TecnaiG^2^ Spirit BioTWIN and images were recorded with an AMT 2k CCD camera.

### *S. aureus* overexpression constructs and minimal inhibitory concentration determination

Fatty acid biosynthesis genes were PCR-amplified using the primers listed in Table S4. PCR products were inserted into pOS1-P*lgt* plasmid and transformed using NEBuilder® HiFi DNA Assembly Cloning Kit (12). *E. coli* transformants were selected for ampicillin (100 *μ*g/ml) resistance, plasmids isolated, and transformed into *S. aureus* RN4220 by electroporation. *fabHB* C113A mutant construct was generated from wiltype pOS1-P*lgt fabHB* with Q5® Site-Directed Mutagenesis Kit (NEB, Ipswich, MA). Chloramphenicol supplemented at 10 *μ*g/ml was used to select for transformants and to maintain the plasmid in *S. aureus*. Approximately 105 CFUs were inoculated into 1 ml liquid media containing two-fold dilutions of amycomicin or platensimycin and incubated with shaking overnight in 13 × 100 mm glass test tubes. The lowest dilution of amycomicin or platensimycin preventing visible growth of *S. aureus* was recorded as the minimal inhibitory concentration for that strain.

### Amycomicin activity against diverse species

Amycomicin solution was spotted onto 90 mm petri dishes containing 15 ml 1X PTSYM pH6.0 or tryptic soy agar pH6.0 (for *Streptococcus pyogenes)*. Once absorbed the following bacterial species were spread on the plates with cotton-tipped applicators: *Macrococcus caseolyticus* ATCC 13518, *Enterococcus faecalis* V583, *Streptococcus pyogenes* 854, *Mycobacterium smegmatis* MC2 155, *Streptomyces coelicolor* A3(2)/M145, *Pseudomonas aeruginosa* PAO1, *Escherichia coli* DH5*α*, and *Candida albicans* SC5314. The petri dishes were incubated at 30 °C.

#### B. subtilis fabHA and fabHB mutants

Mutations in *fabHA* and *fabHB* were generated in *B. subtilis* 168 genome as described previously (13). Genomic DNA for mutagenesis was kindly provided by David Rudner (Harvard Medical School). Amycomicin solution was spotted onto 100 mm petri dishes containing 15 ml 1X PTSYM pH6.0. Once absorbed the bacteria were spread on the plates with cotton-tipped applicators and incubated at 22 °C for 48 hours.

### Murine skin infections

Six to seven-week-old female BALB/Cj mice were infected using a tape-stripping model of skin infection (14). Approximately 107 CFU of SAP149, a USA300-0114 CA-MRSA strain with *luxBADCE* integrated at USA300HOU_1102 were suspended in 5 *μ*l phosphate-buffered saline and inoculated onto the wound (15). Immediately after infection the mice were visualized using IVIS spectrum *in vivo* imaging system with data being collected at 60-second exposure times. Radiance was quantified for individual mice using Living Image Version 4.1.0.11858 software (Caliper LifeSciences) software. Upon acquisition of the images, circles of the same area were drawn around the infection sites and elsewhere in the frame. Background radiance acquired from areas outside of infection was subtracted from radiance emanating from the infection sites. The infection proceeded untreated for 24 hours and luminescence was measured again prior to application of the first treatment. The mice were separated into three groups, receiving 0.001% amycomicin, 2% mupirocin ointment (TARO pharmaceuticals, Hawthorne, NY), or no treatment. Twenty microliters of amycomicin were applied with a pipette tip to the wound, while mupirocin ointment was applied liberally with a cotton-tipped applicator. The treatment was applied 3 times total at 24-hour intervals. Bioluminescence data were acquired daily as well. Twenty-four hours after the last treatment (day 4 post-infection and 3 days after the first treatment) the last luminescence data point was acquired. The skin samples were homogenized using Bullet Blender (Next Advance, Troy, NY), serially diluted and plated for determination of CFU/ml. Throughout the experiment the mice were handled according to Vanderbilt University IACUC regulations.

---

**Fig. S1:**
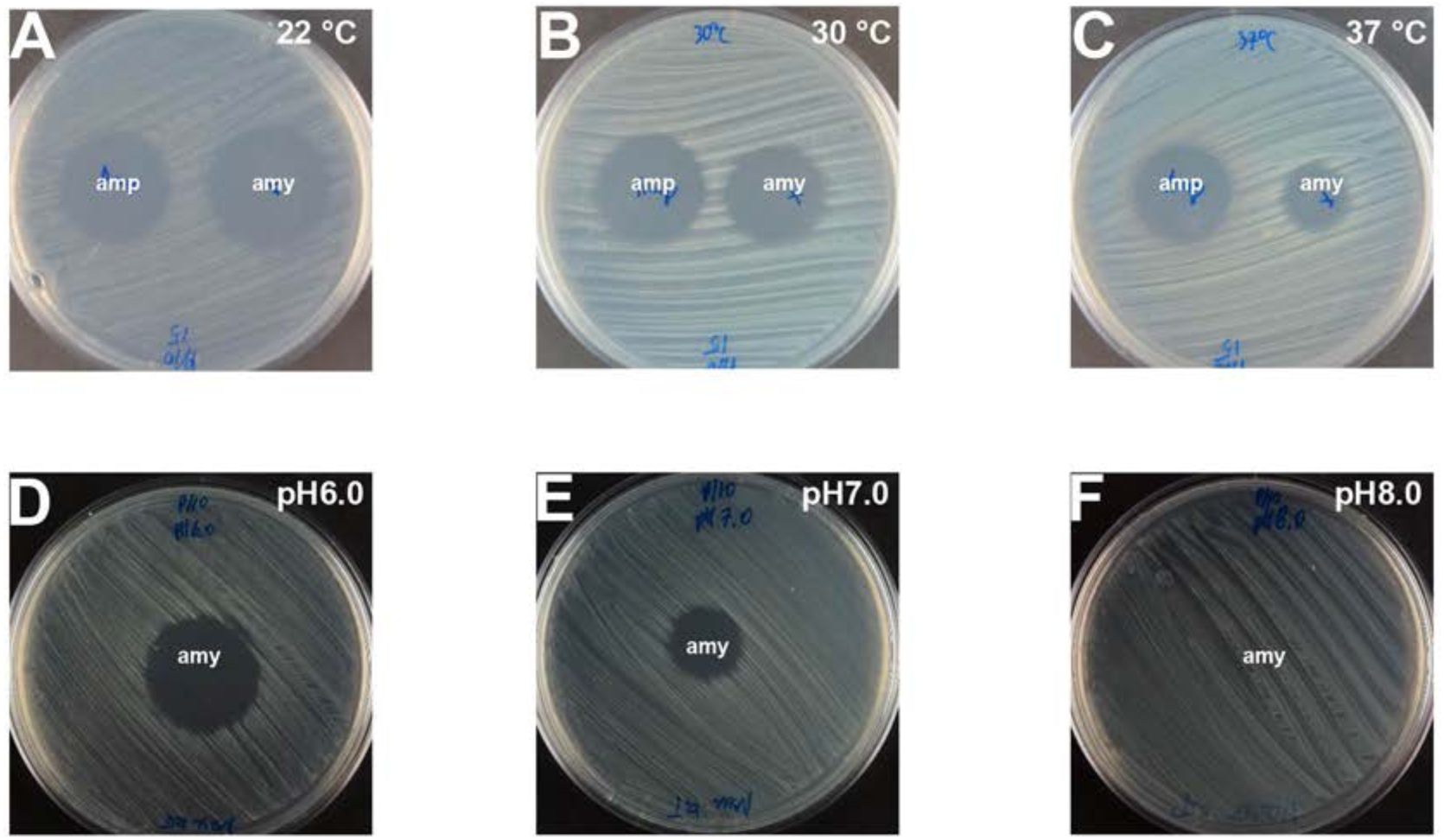
Temperature and pH dependence of amycomicin activity. Zones of inhibition produced by 1 *μ*g ampicillin (Amp) and 100 ng amycomicin (Amy) on lawns of *S. aureus* grown at indicated temperatures (*A-C*) and pH (*D-F*).

**Fig. S2:**
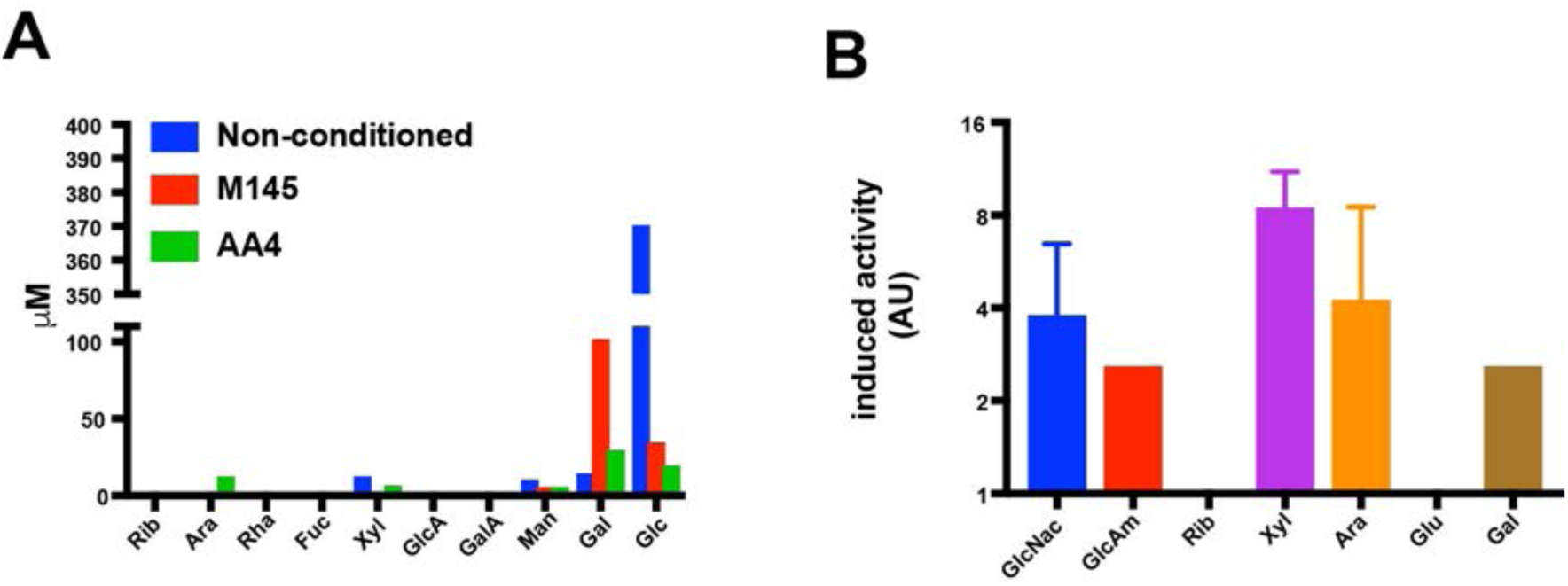
Carbohydrates induce production of antimicrobial activity by AA4. (*A*) Concentration of individual saccharides detected in fresh (non-conditioned) medium, medium conditioned by M145 and medium conditioned by AA4. (*B*) Amount of antimicrobial activity expressed in arbitrary units (AU) produced by AA4 and induced by 1mM of the indicated carbohydrate (see Materials and Methods). Abbreviations: Rib=ribose, Ara=arabinose, Rha=rhamnose, Fuc=fucose, Xyl=xylose, GlcA=glucuronic acid, GalA=galacturonic acid, Man=mannose, Gal=galactose, Glc=glucose, Glc*N*ac=*N*-acetyl-glucosamine, GlcAm=glucosamine.

## Analytical Data for Amycomicin (1)

**Fig. S3A:**
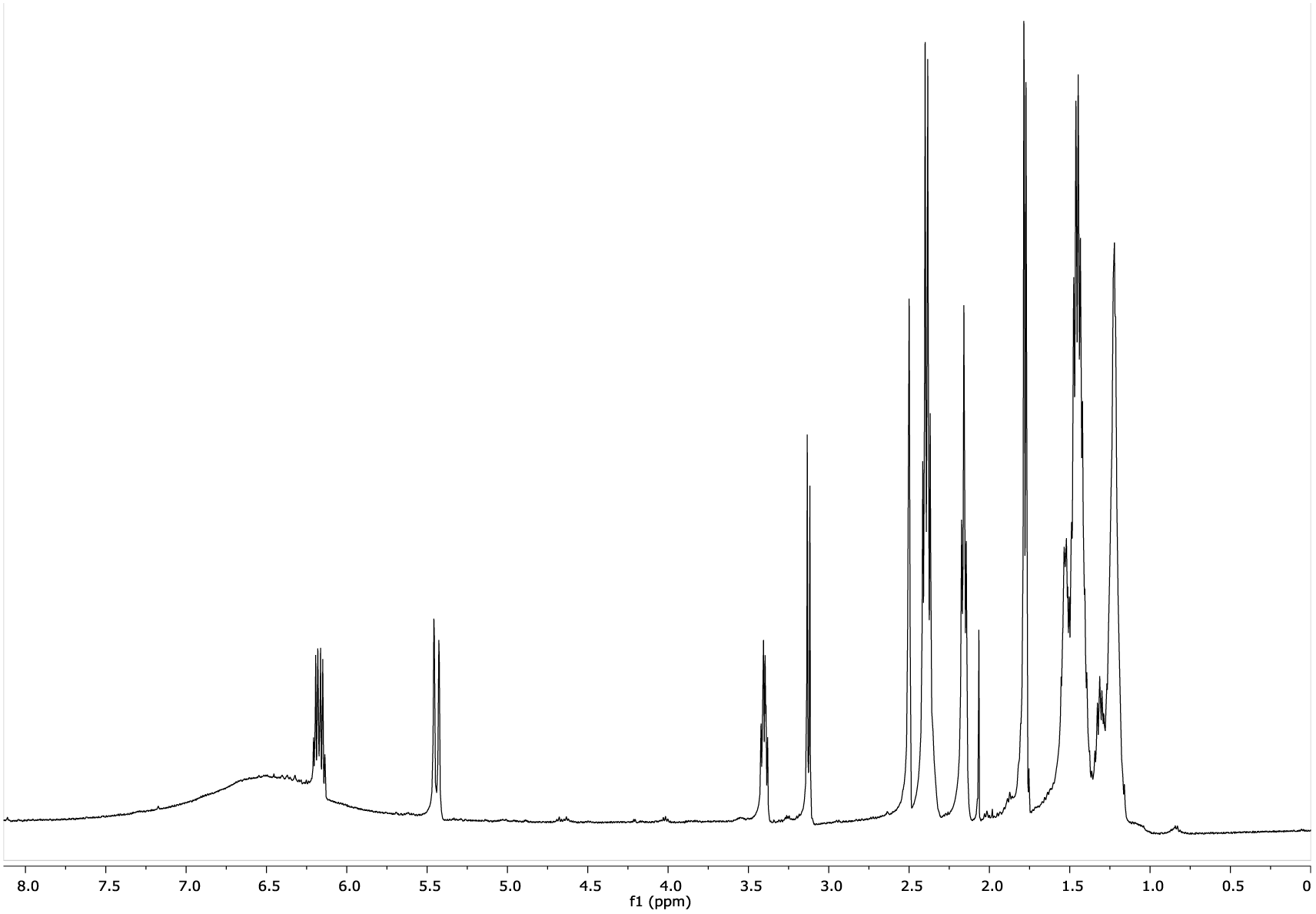
^1^H NMR spectrum of amycomicin (1) in *d_6_*-DMSO (500 MHz)

**Fig. S3B:**
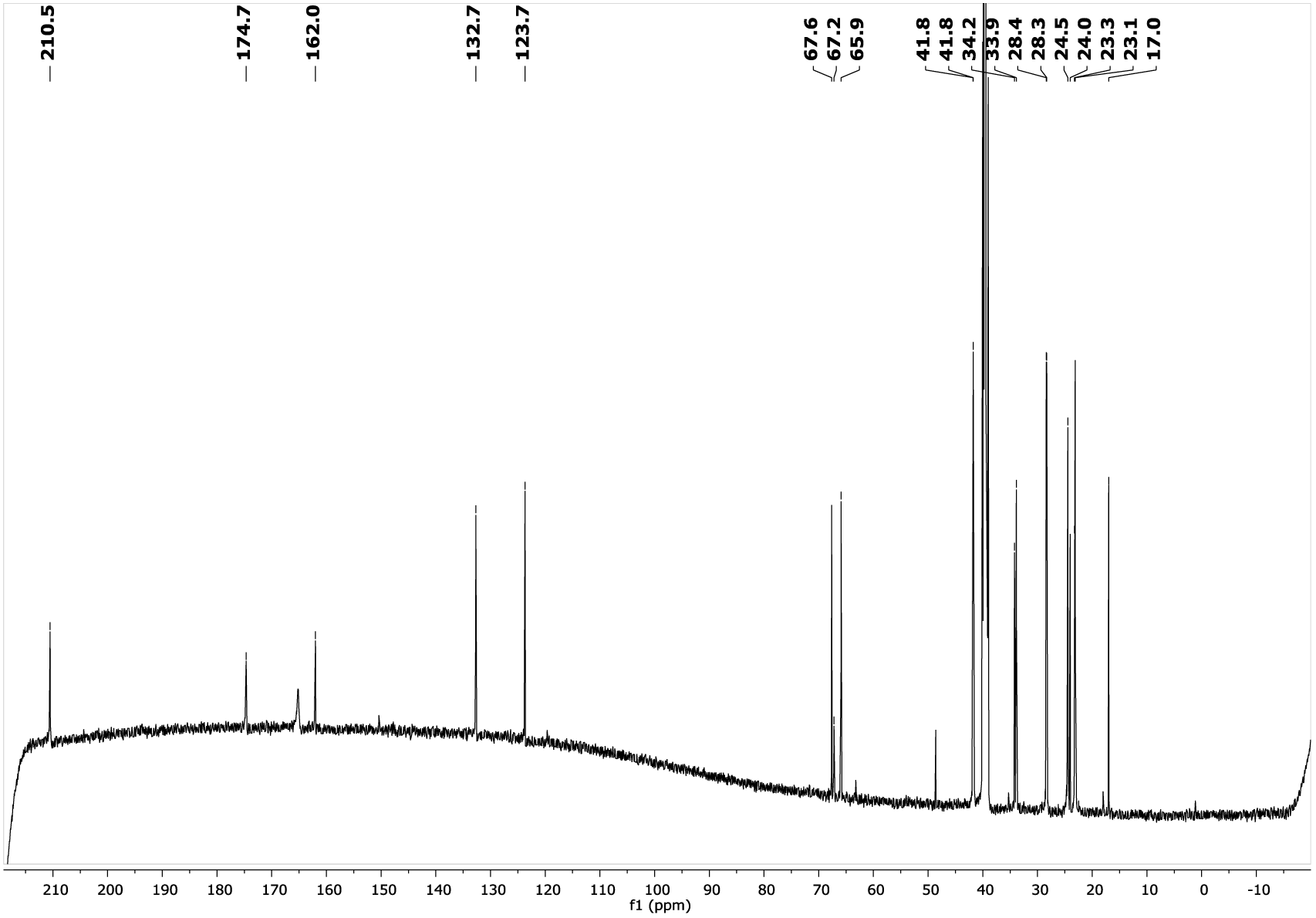
^13^C NMR spectrum of amycomicin (1) in *d_6_*-DMSO (125 MHz)

**Fig. S3C:**
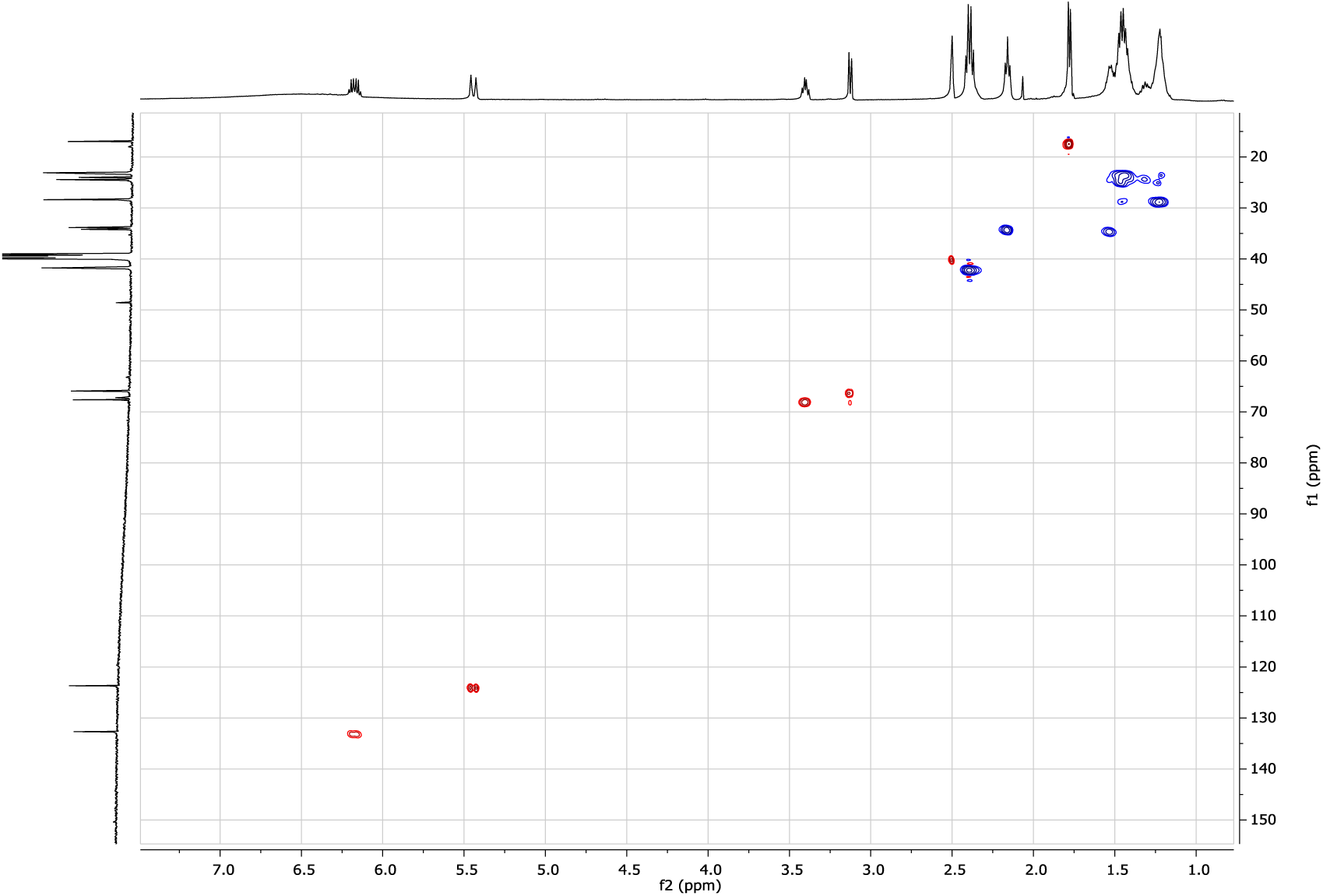
gHSQC spectrum of amycomicin (1) in d_6_-DMSO (^1^H - 500 MHz)

**Fig. S3D:**
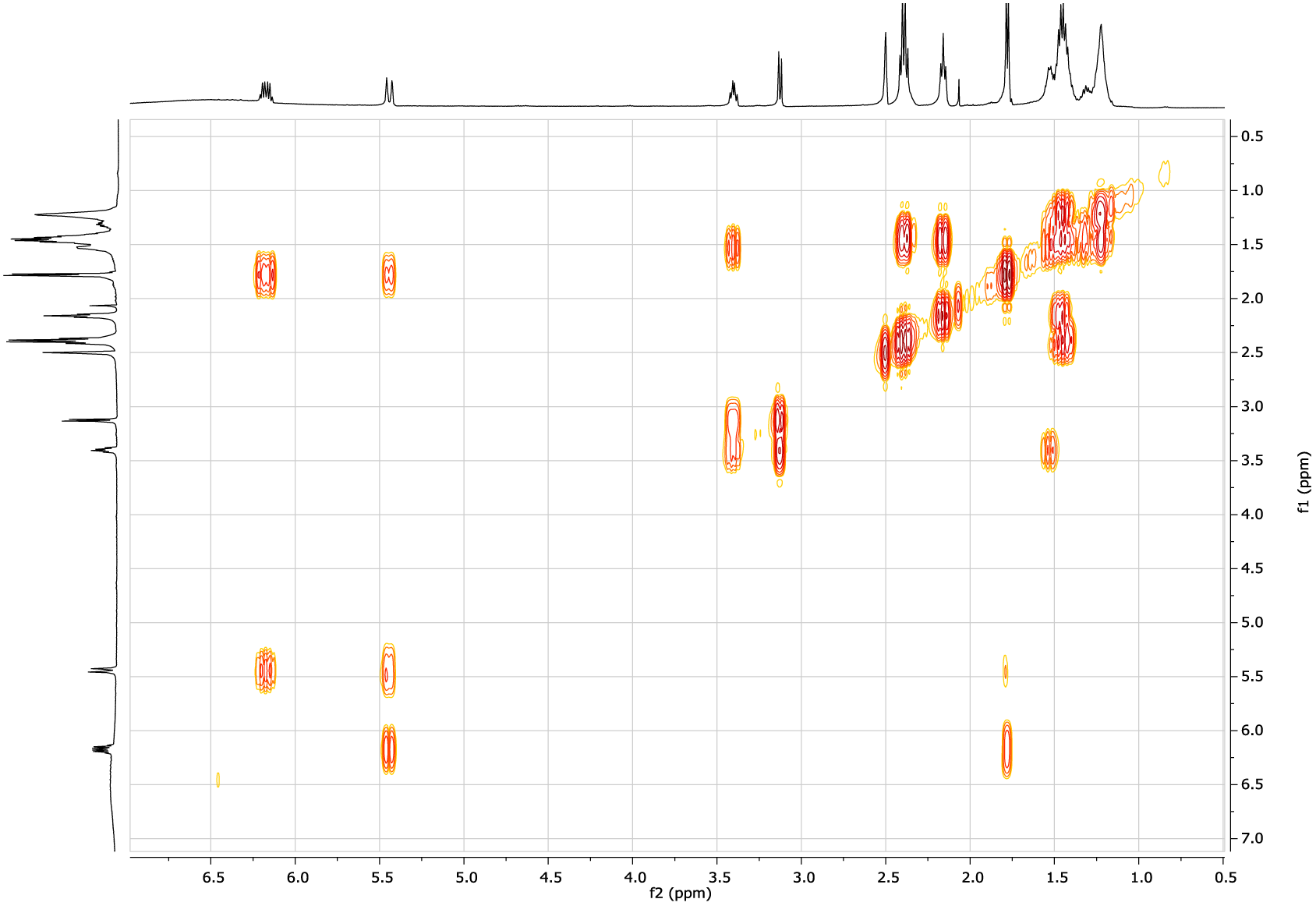
dqfCOSY spectrum of amycomicin (1) in d_6_-DMSO (500 MHz)

**Fig. S3E:**
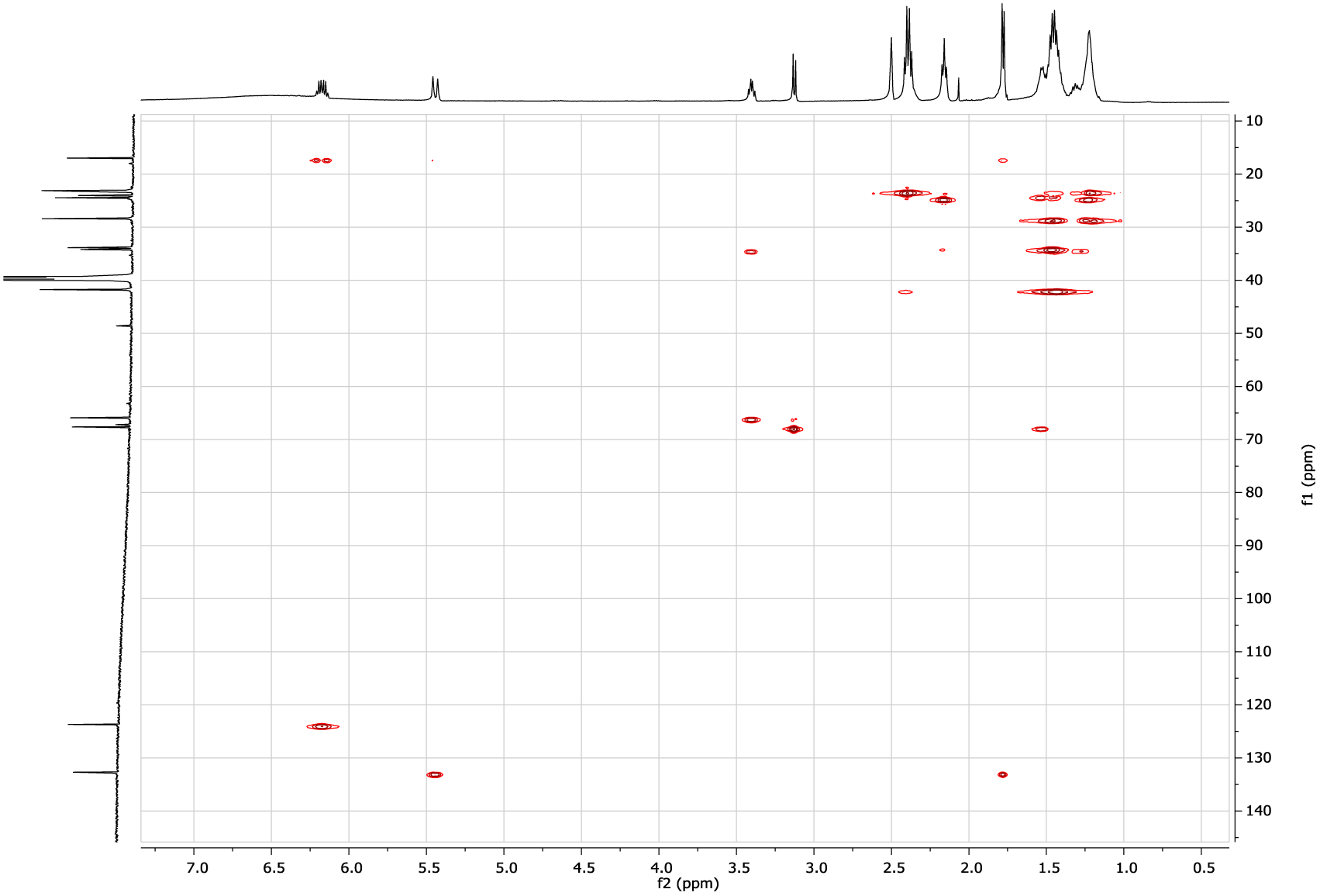
H2BC spectrum of amycomicin (1) in *d_6_*-DMSO (^1^H - 500 MHz)

**Fig. S3F:**
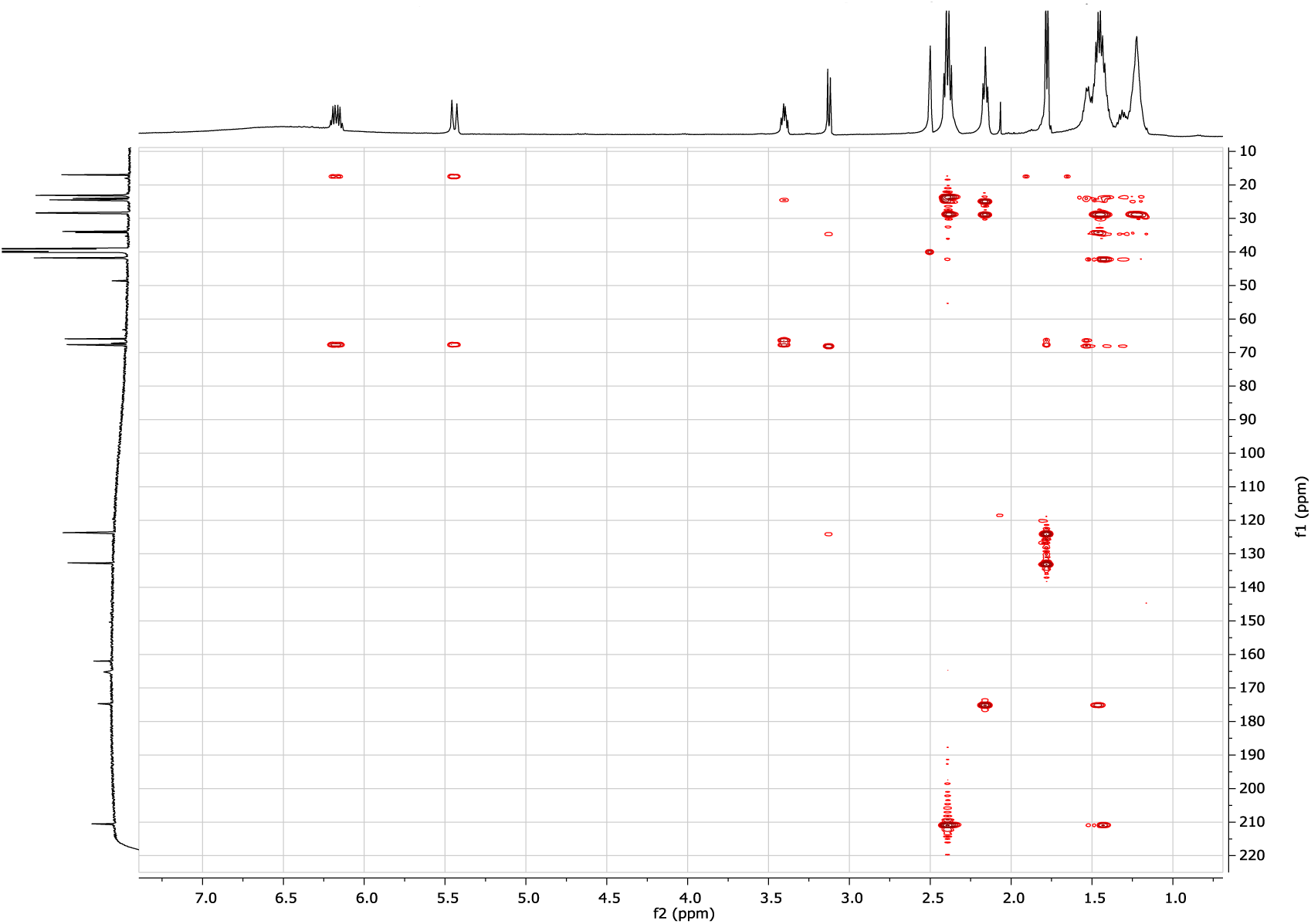
gHMBCAD spectrum of amycomicin (1) in *d_6_*-DMSO (^1^H - 500 MHz)

**Fig. S3G:**
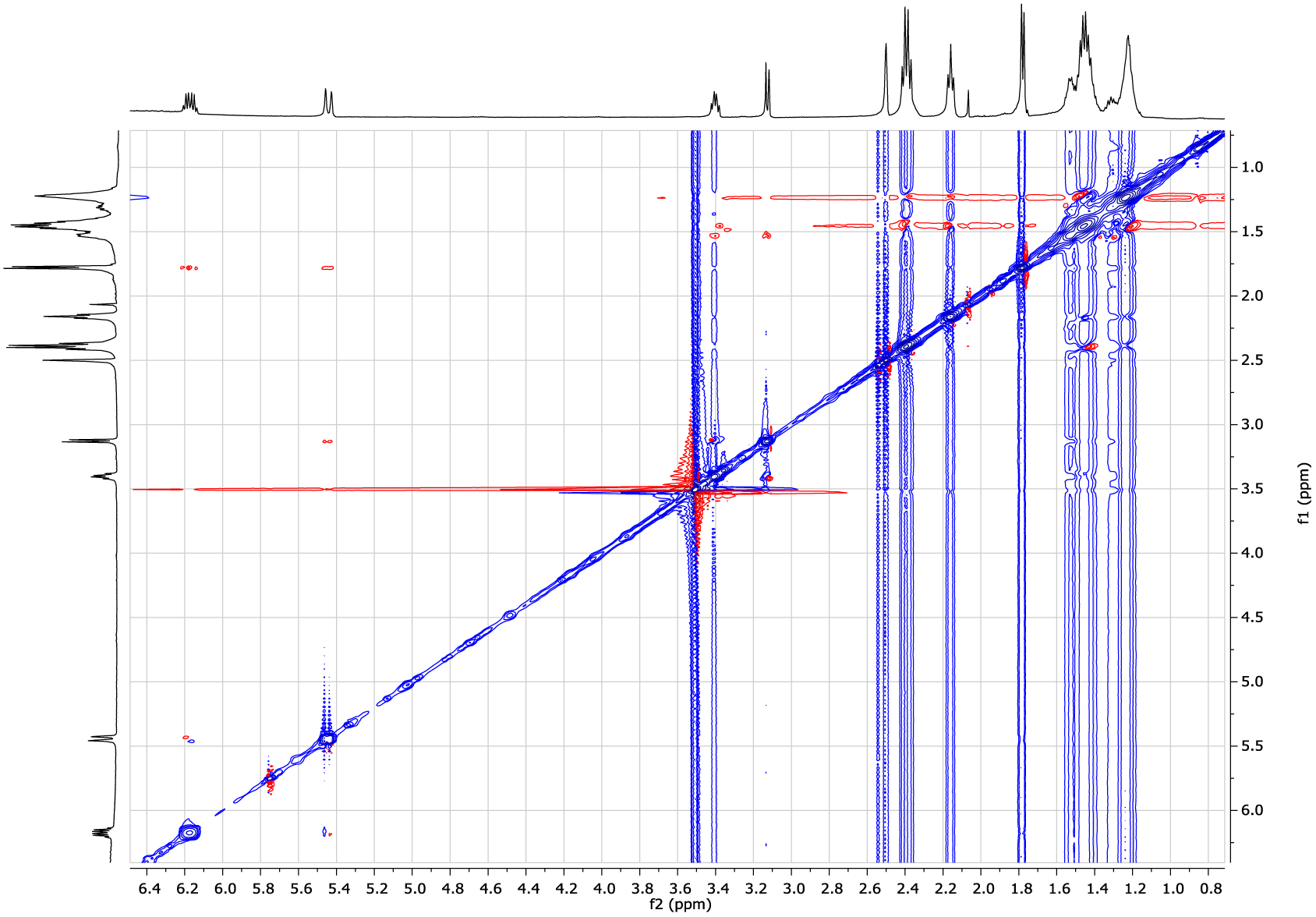
ROESY spectrum of amycomicin (1) in *d_6_*-DMSO (500 MHz)

**Table S1.** NMR assignments for amycomicin (1) in *d_6_*-DMSO.

| position | $\delta_C^b$ | $\delta_H$ (*J* in Hz)<sup>a</sup> | COSY<sup>a</sup> | HMBC<sup>c</sup> | H2BC | ROESY |
| --- | --- | --- | --- | --- | --- | --- |
| 1 | 174.7 |  |  |  |  |  |
| 2 | 33.9 | 2.16 (t; 7.3) | 3 | 1, 3, 4 | 3 |  |
| 3 | 22.2 | 1.47 (m) |  | 1, 2, 4 | 2, 4 |  |
| 4 | 28.4 | 1.23 (m) |  | 2, 3, 5 | 3, 5 |  |
| 5 | 28.3 | 1.23 (m) |  | 4, 6, 7 | 4, 6 |  |
| 6 | 23.1 | 1.44 (m) | 7 | 5, 7, 8 | 5, 7 |  |
| 7 | 41.75 | 2.40 (t; 7.3) | 6 | 5, 6, 8 | 6 |  |
| 8 | 210.5 |  |  |  |  |  |
| 9 | 41.77 | 2.38 (t; 7.3) | 10 | 8, 10 | 10 |  |
| 10 | 23.2 | 1.44 (m) | 9 | 8, 9, 11, 12 | 9 |  |
| 11 | 24.0 | 1.32 (m) |  | 9, 10, 12 | 10, 12 |  |
| 12 | 34.2 | 1.53 (m) | 13 | 11, 13, 14 | 11, 13 | 13, 14 |
| 13 | 67.6 | 3.41 (dt; 7.9, 4.9) | 12, 14 | 11, 12, 14, 15 | 12, 14 | 12, 14 |
| 14 | 65.9 | 3.13 (d; 7.8) | 13 | 12, 13, 16 | 13 | 12, 13, 16 |
| 15 | 67.2 |  |  |  |  |  |
| 16 | 123.7 | 5.46 (d; 15.1) | 17, 18 | 15, 18 | 17 | 14, 17, 18 |
| 17 | 132.7 | 6.19 (dq; 15.1, 6.8) | 16, 18 | 15, 16, 18 | 16, 18 | 16, 18 |
| 18 | 17.0 | 1.78 (d; 6.8) | 16, 17 | 16, 17 | 17 | 16, 17 |
| 19 | 162.0 |  |  |  |  |  |

<sup>a</sup> 500 MHz for <sup>1</sup>H NMR, dqfCOSY, gHSQC, HMBC, and ROESY<sup>b</sup> 125 MHz for <sup>13</sup>C NMR

**Fig. S3H:**
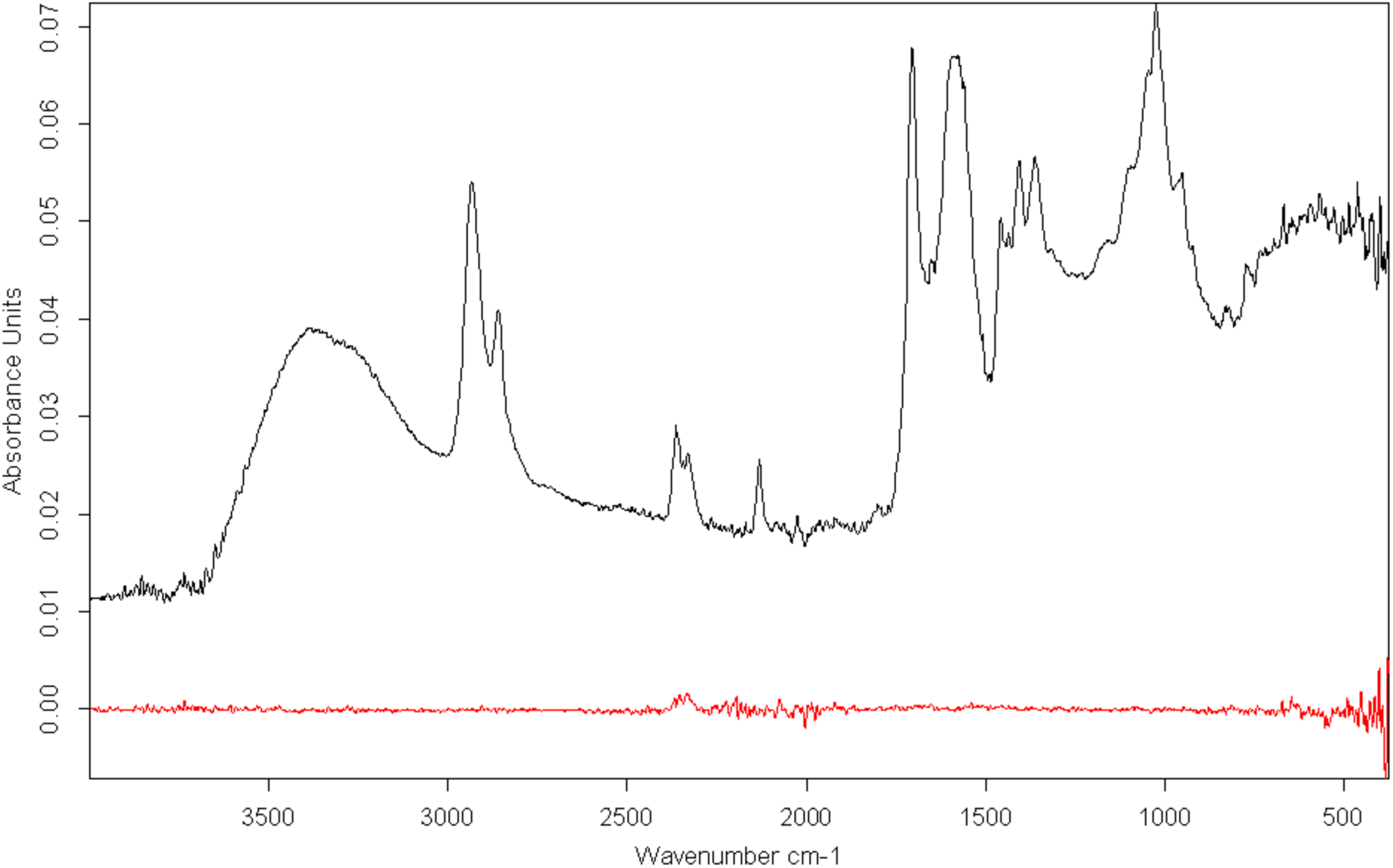
IR spectrum of amycomicin (1) (neat)

## Analytical Data on Epoxyketone Amycomicin (2)

**Fig. S4A:**
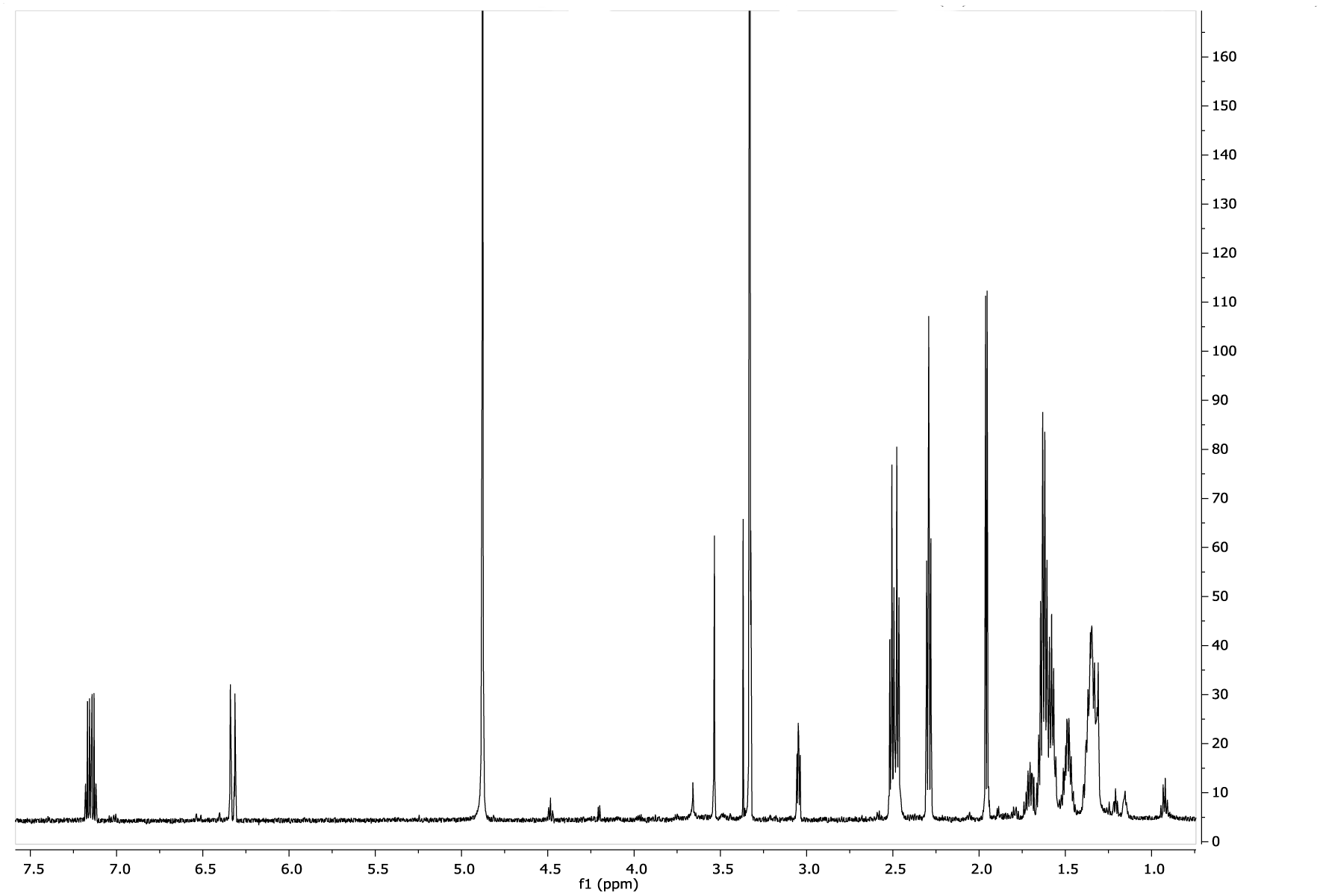
^1^H NMR spectrum of epoxyketone amycomicin (2) in CD_3_OD (600 MHz)

**Fig. S4B:**
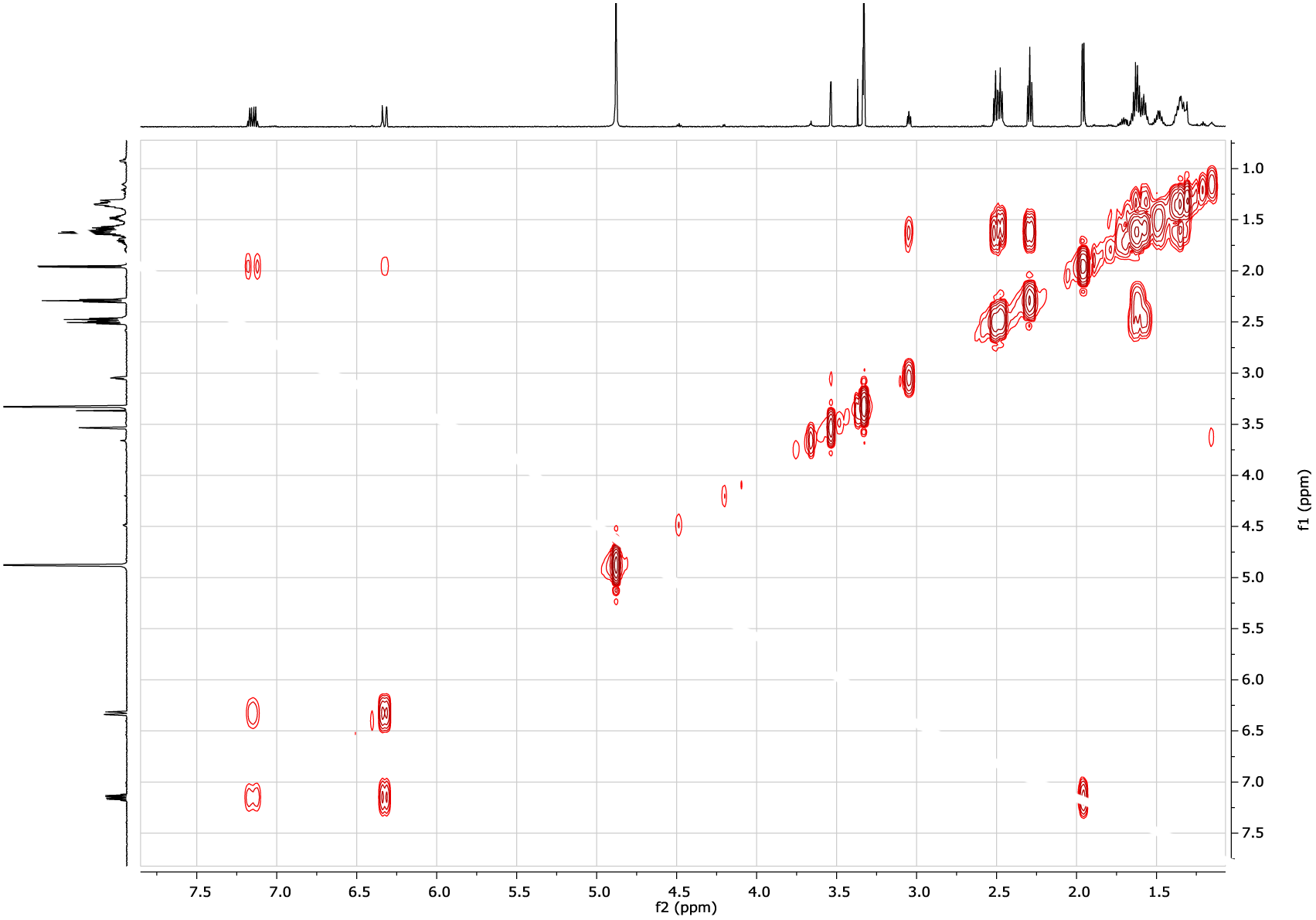
dqfCOSY spectrum of epoxyketone amycomicin (2) in CD_3_OD (600 MHz)

**Fig. S4C:**
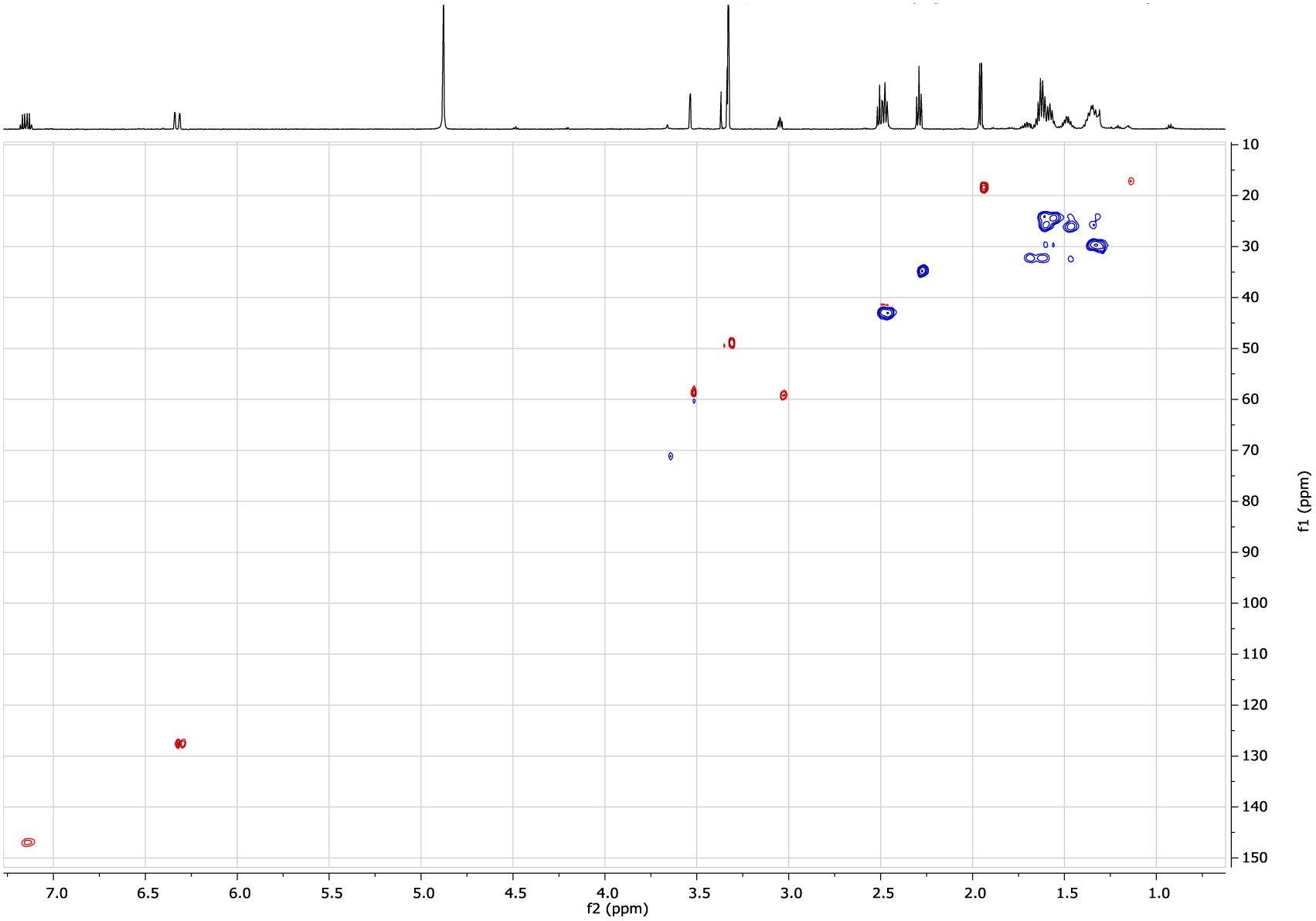
gHSQC spectrum of epoxyketone amycomicin (2) in CD_3_OD (1H - 600 MHz)

**Fig. S4D:**
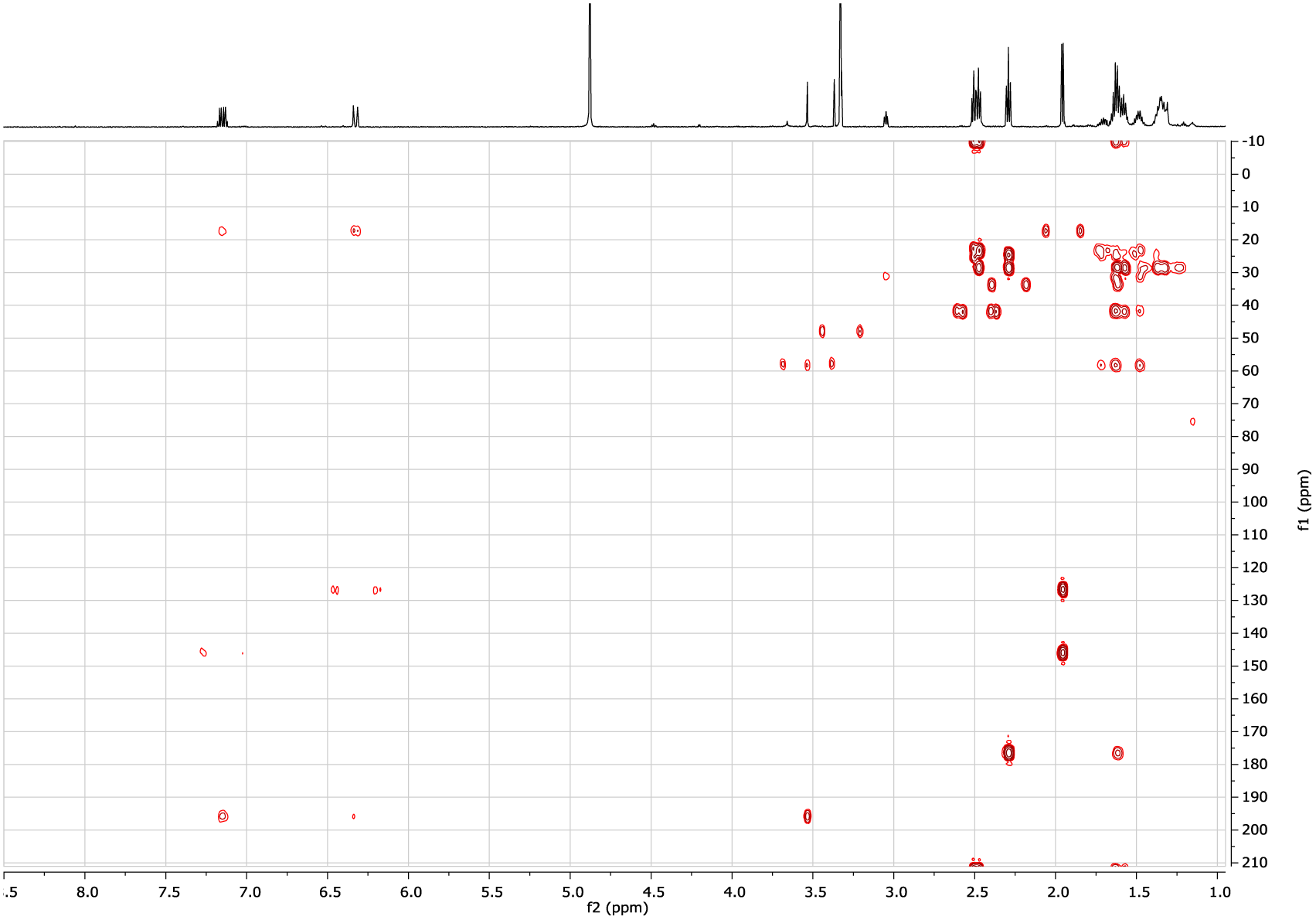
gHMBCAD spectrum of epoxyketone amycomicin (2) in CD_3_OD (^1^H - 600 MHz)

**Fig. S4E:**
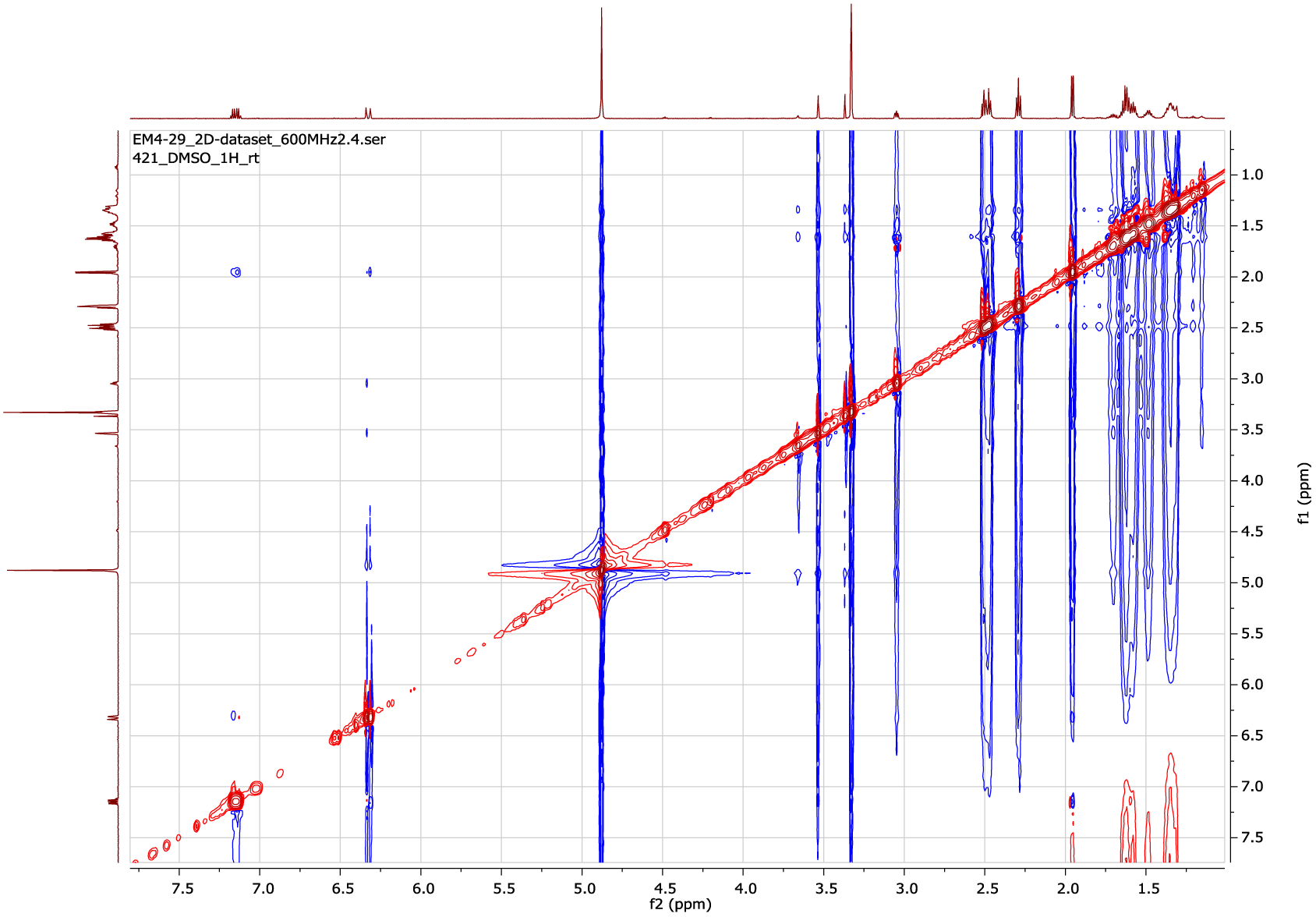
ROESY spectrum of epoxyketone amycomicin (2) in CD_3_OD (600 MHz)

**Table S2.** NMR assignments for epoxyketone amycomicin (2) in CD_3_OD.

| position | $\delta_C^b$ | $\delta_H$ (*J* in Hz)<sup>a</sup> | COSY<sup>a</sup> | HMBC<sup>c</sup> |
| --- | --- | --- | --- | --- |
| 1 | 177.0 |  |  |  |
| 2 | 34.8 | 2.27 (t; 7.4) | 3 | 1, 4 |
| 3 | 25.6 | 1.60 (m) | 2 | 1, 2, 4 |
| 4 | 29.5 | 1.34 (m) |  | 3, 5 |
| 5 | 29.9 | 1.31 (m) |  | 4 |
| 6 | 24.4 | 1.56 (m) | 7 | 5, 8 |
| 7 | 43.1 | 2.45 (t; 7.4) | 6 | 5, 6, 8 |
| 8 | 212.9 |  |  |  |
| 9 | 42.8 | 2.49 (t; 7.2) | 10 | 8, 10, 11 |
| 10 | 24.1 | 1.61 (m) | 9 | 8, 9, 12 |
| 11 | 26.1 | 1.46 (m) |  | 10, 12, 13 |
| 12a | 32.3 | 1.69 (m) | 13 | 11, 13 |
| 12b |  | 1.62 (m) | 13 | 11, 13 |
| 13 | 59.2 | 3.03 (ddd; 11.1, 4.9, 2.0) | 12, 13 | 11, 12, 14 |
| 14 | 58.6 | 3.51 (d; 2.0) | 13 | 12, 13, 15 |
| 15 | 196.9 |  |  |  |
| 16 | 127.6 | 6.30 (dq; 15.8, 1.7) | 17, 18 | 15, 18 |
| 17 | 147.1 | 7.14 (dq; 15.8, 6.9) | 16, 18 | 15, 18 |
| 18 | 18.4 | 1.94 (dd; 6.9, 1.7) | 16, 17 | 16, 17 |

<sup>a</sup> 600 MHz for <sup>1</sup>H NMR, dqfCOSY, gHSQC, and HMBC
<sup>b</sup> interpreted from HMBC and gHSQC

**Fig. S4F:**
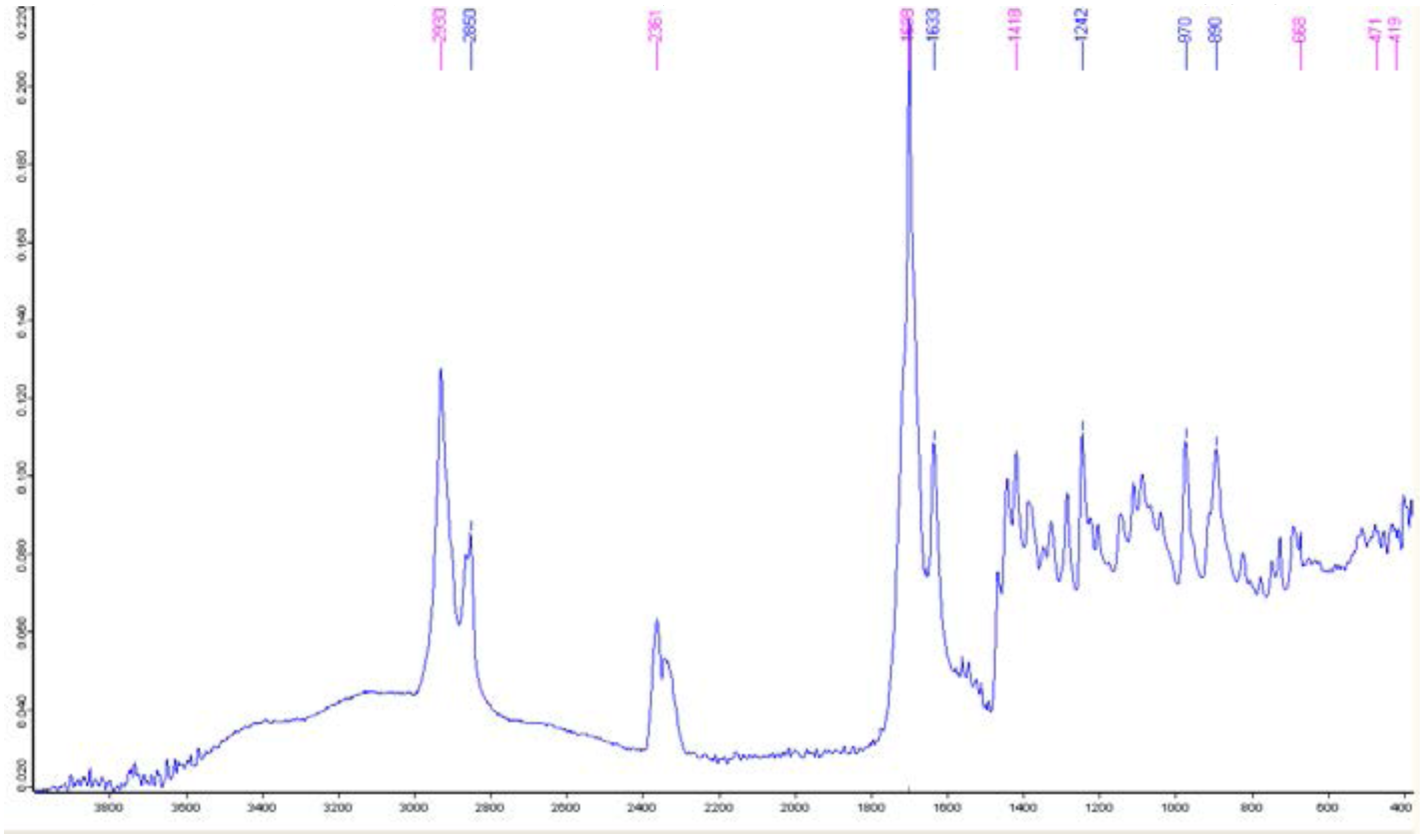
IR spectrum of epoxyketone amycomicin (2) (neat)

**Fig. S4G:**
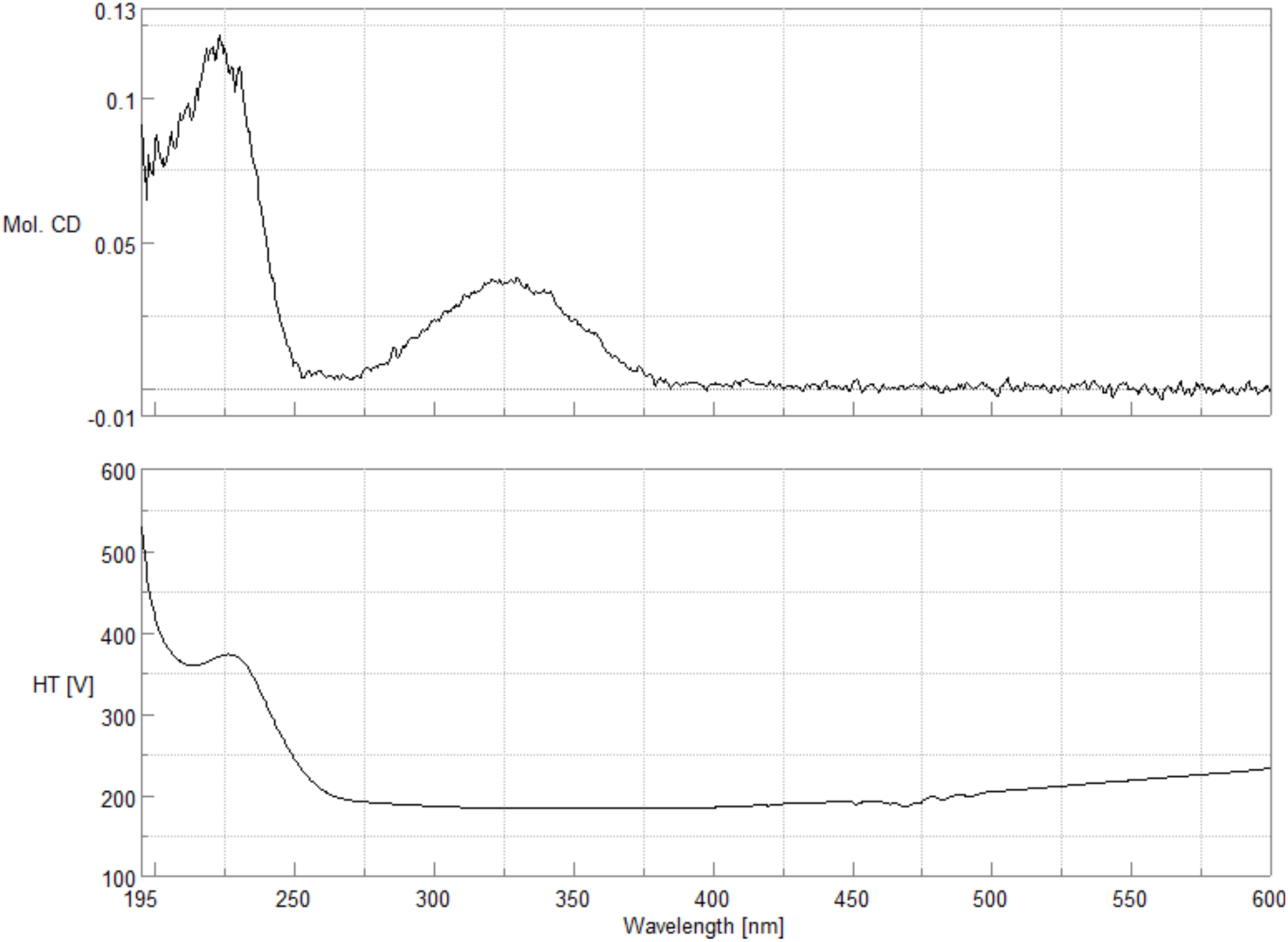
Circular dichroism spectrum on epoxyketone amycomicin (2) in MeOH.

**Fig. S5:**
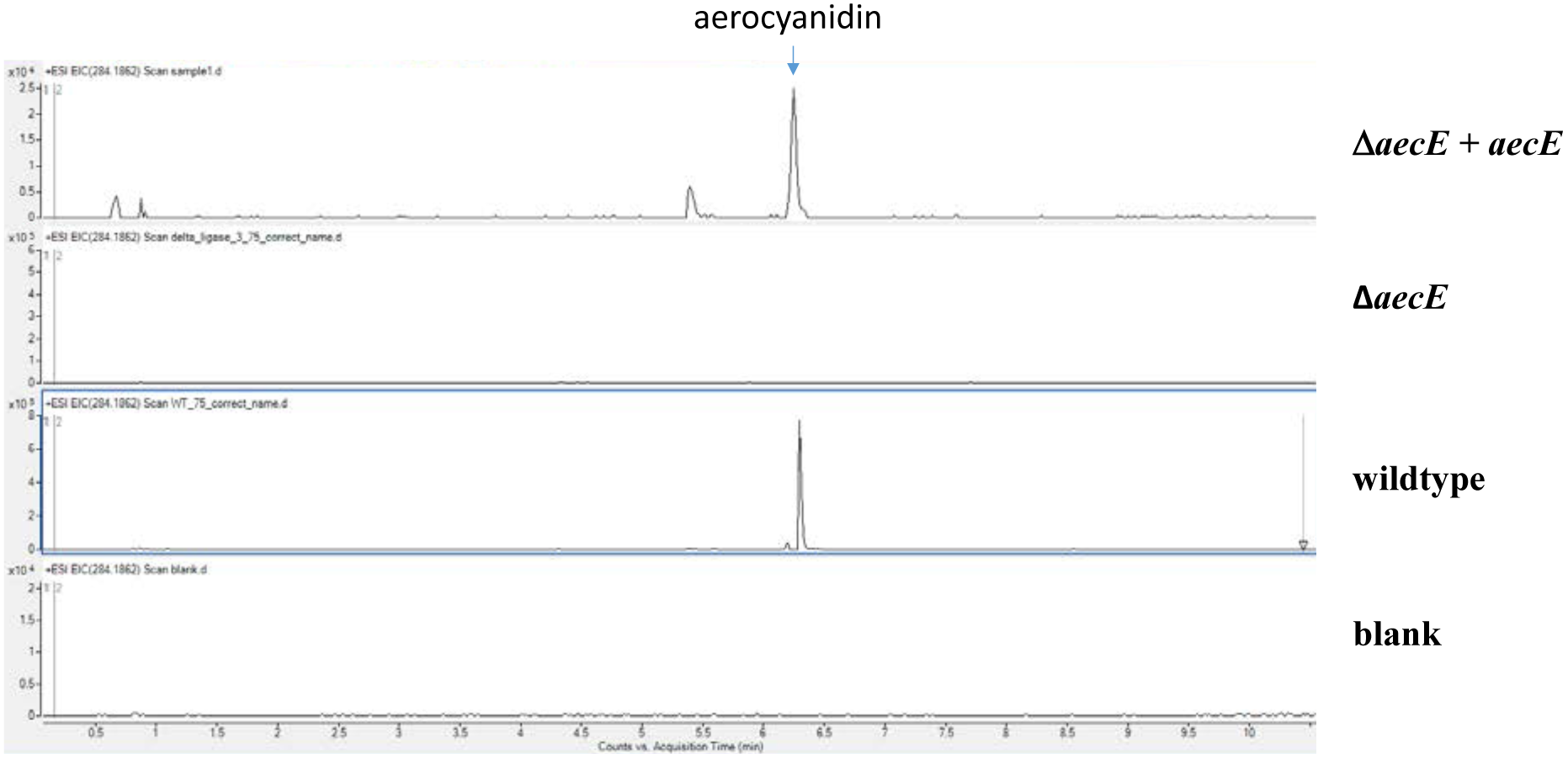
MS analysis of extracted ion chromatogram for production of aerocyanidin.

## Analytical Data on Palmitoleic Acid (3)

**Fig. S6A:**
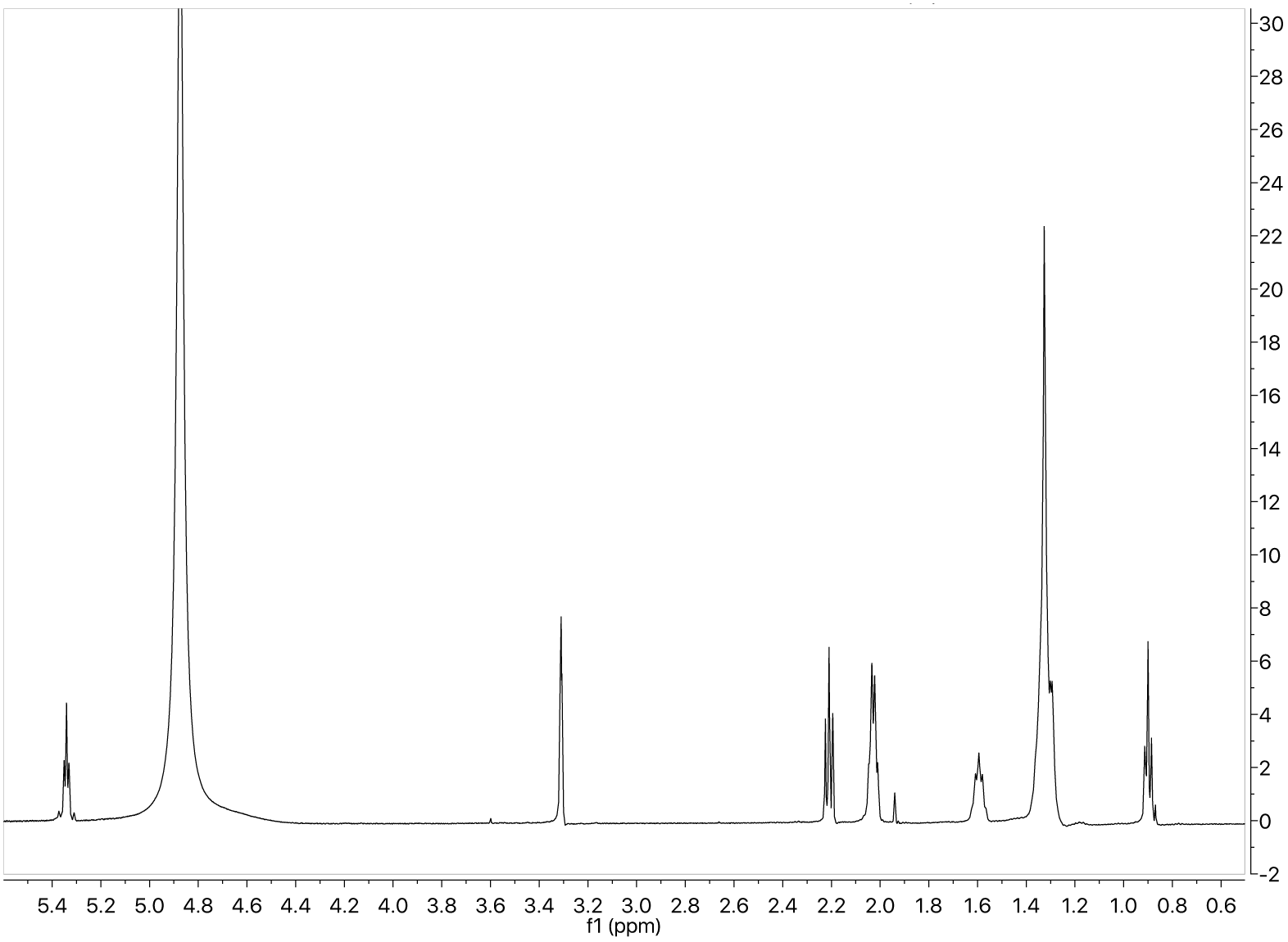
^1^H NMR spectrum of palmitoleic acid (3) in CD_3_OD (600 MHz)

**Fig. S6B:**
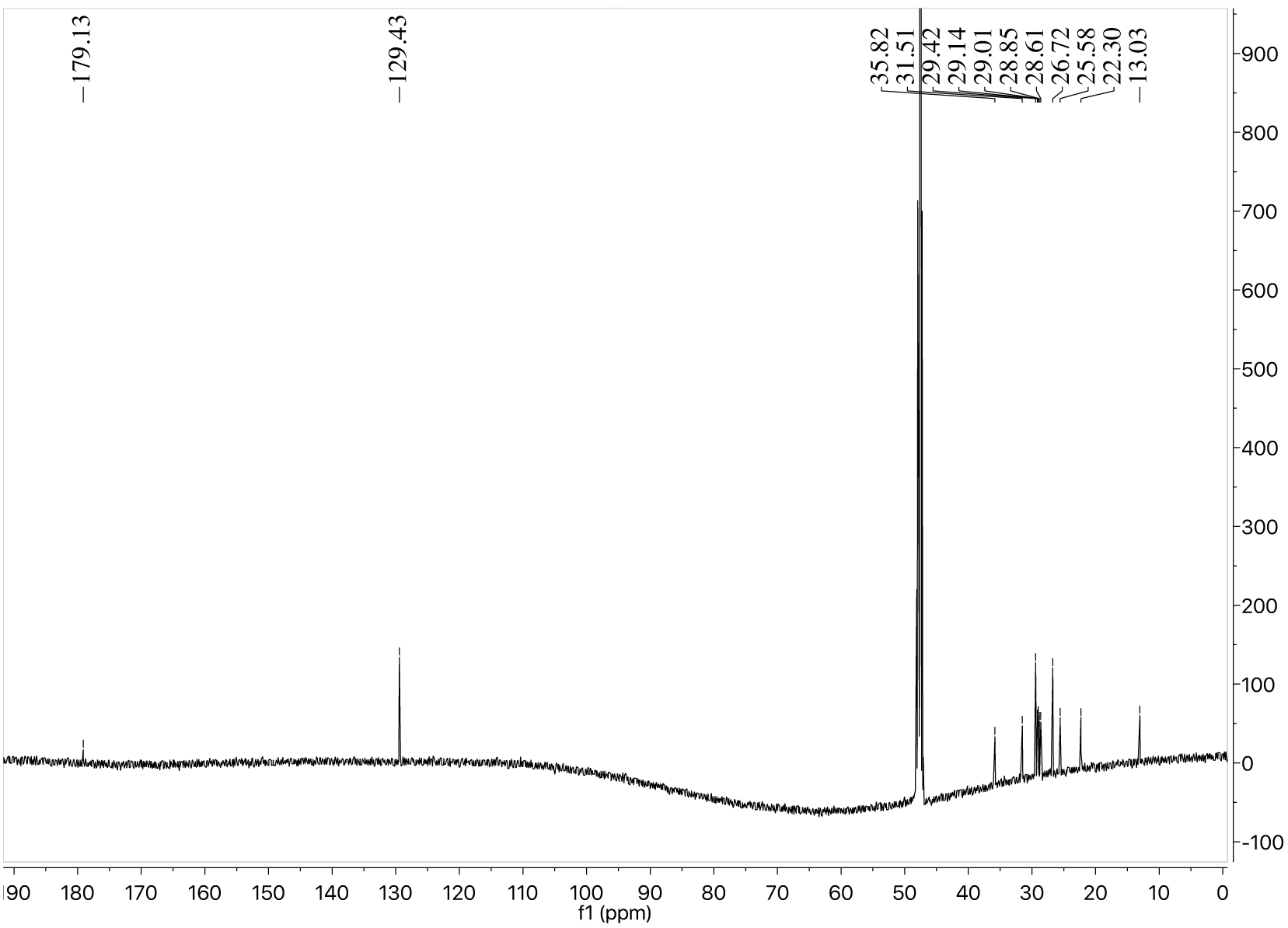
^13^C NMR spectrum of palmitoleic acid (3) in CD_3_OD (125 MHz)

**Fig. S6C:**
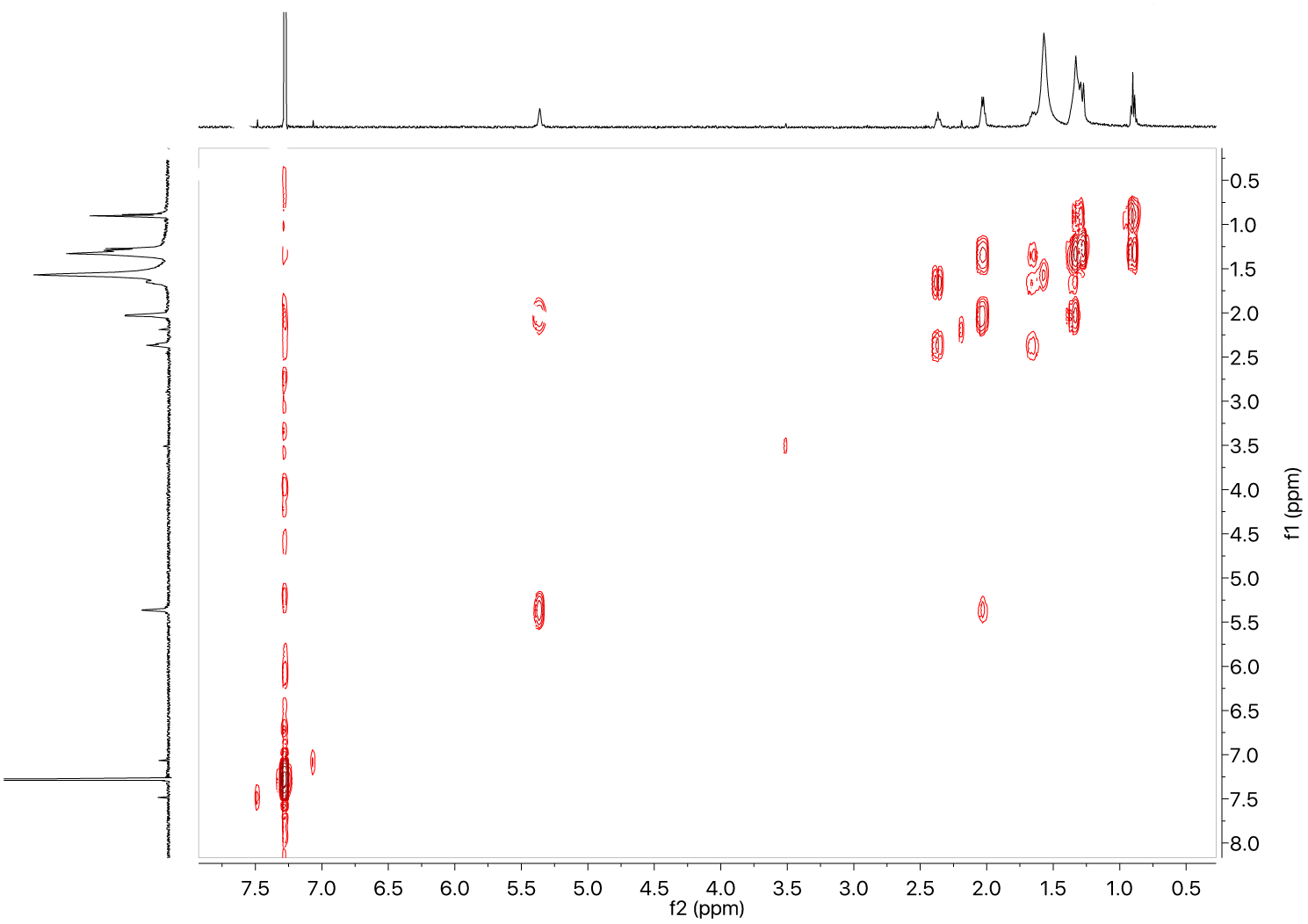
gCOSY spectrum of palmitoleic acid (3) in CDCl_3_ (600 MHz)

**Fig. S6D:**
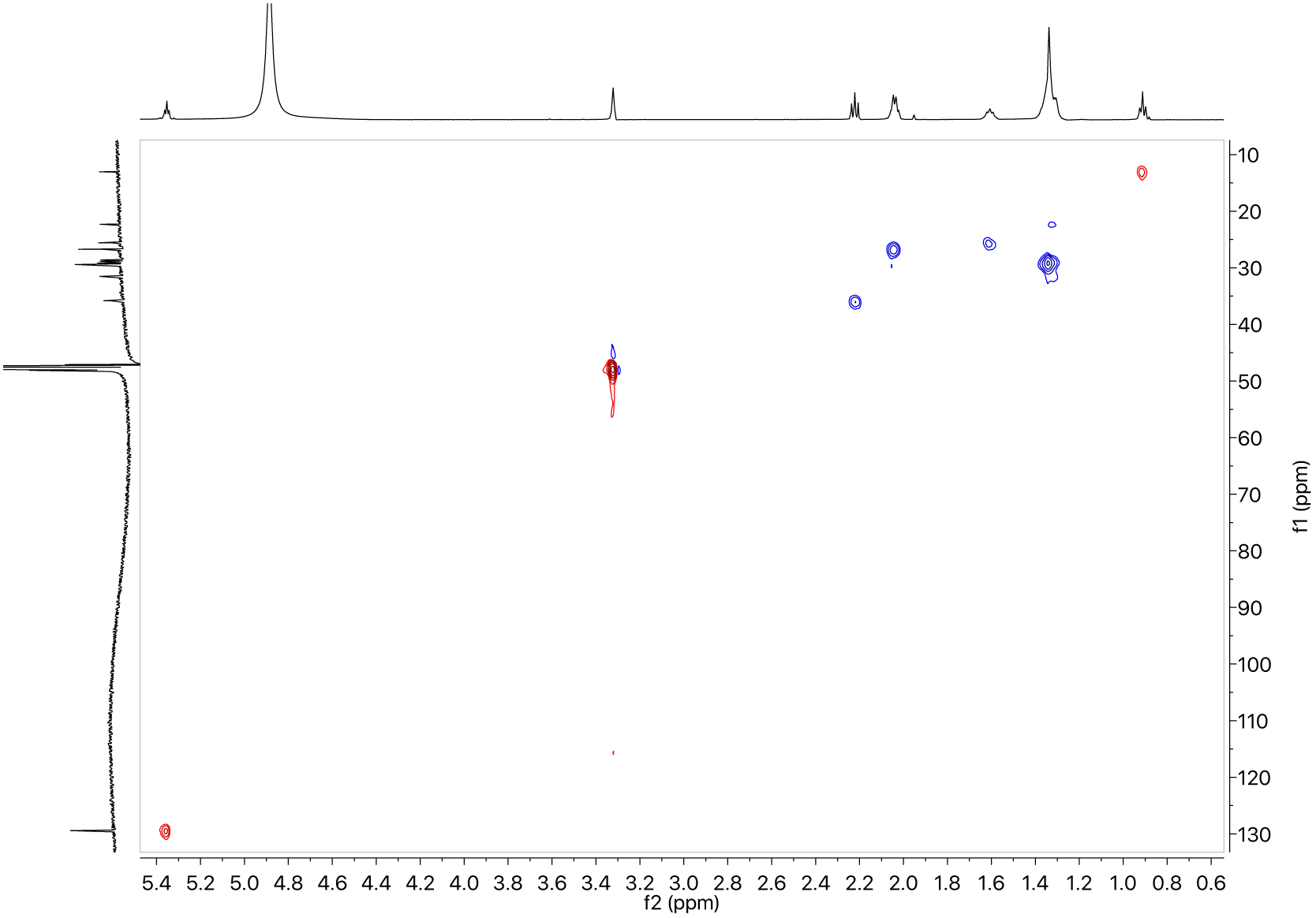
gHSQC spectrum of palmitoleic acid (3) in CD_3_OD (^1^H - 600 MHz)

**Fig. S6E:**
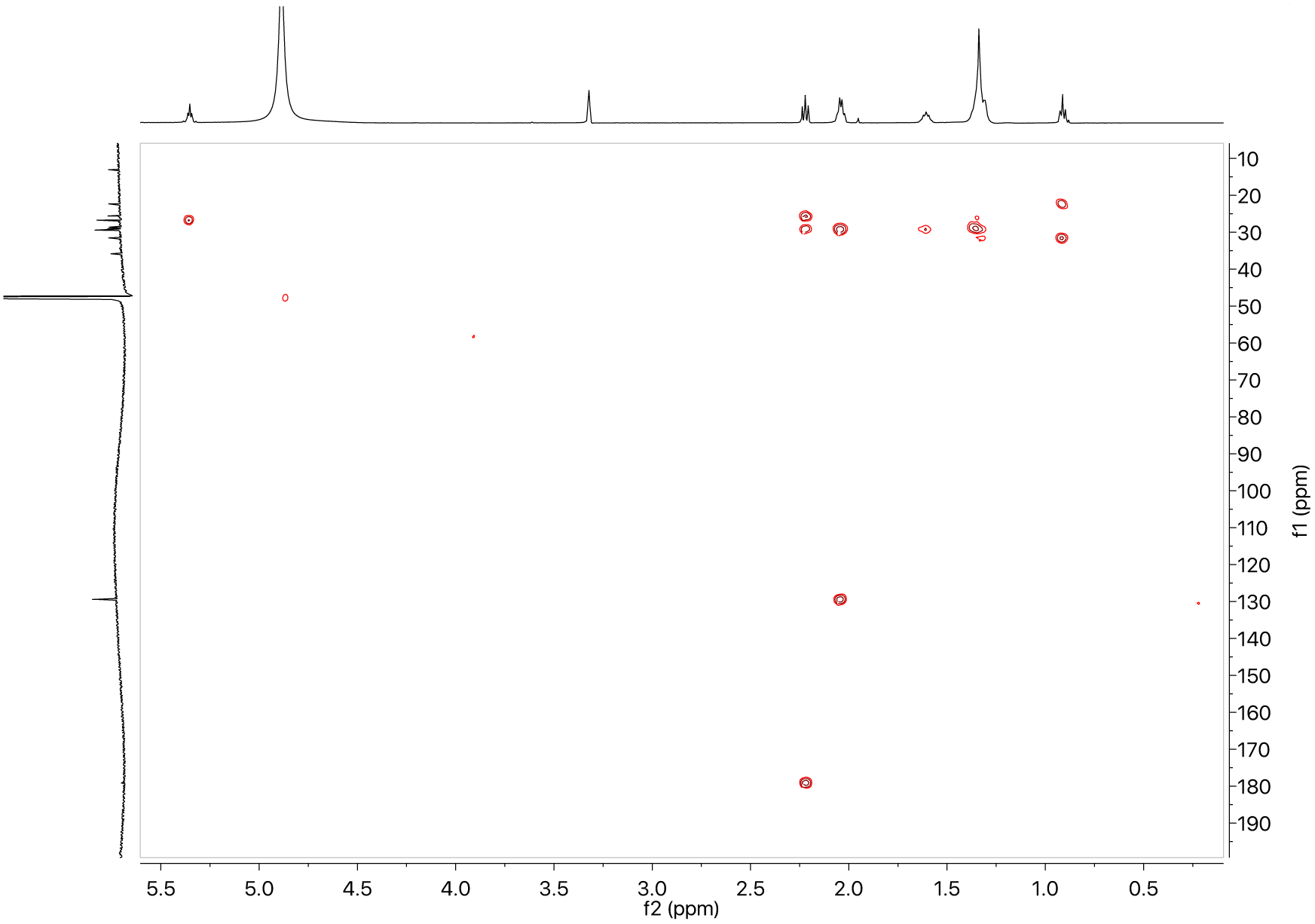
gHMBC spectrum of palmitoleic acid (3) in CD_3_OD (^1^H - 600 MHz)

**Fig. S6F:**
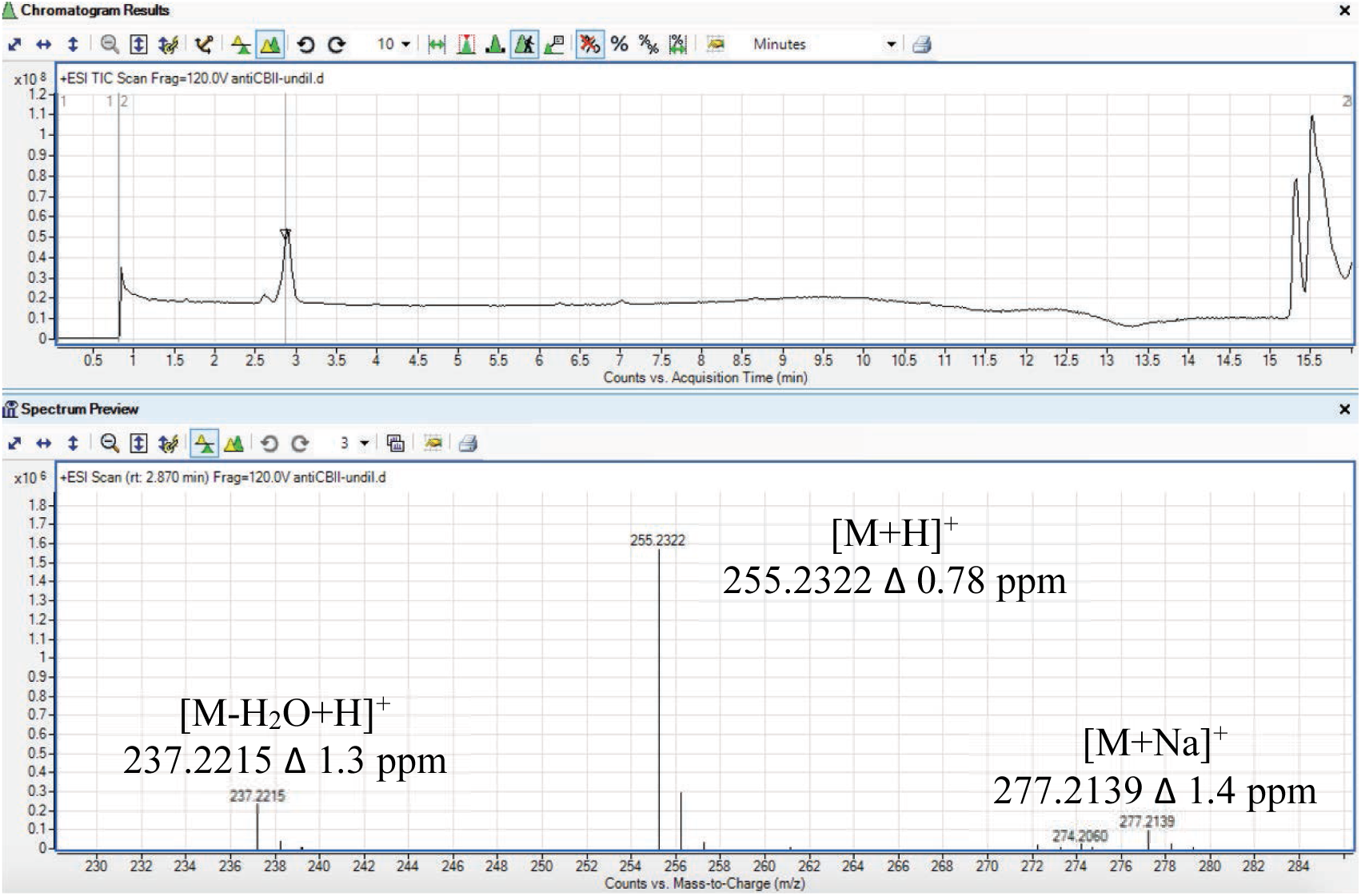
HRLCMS spectrum of palmitoleic acid (3)

**Fig. S6G:**
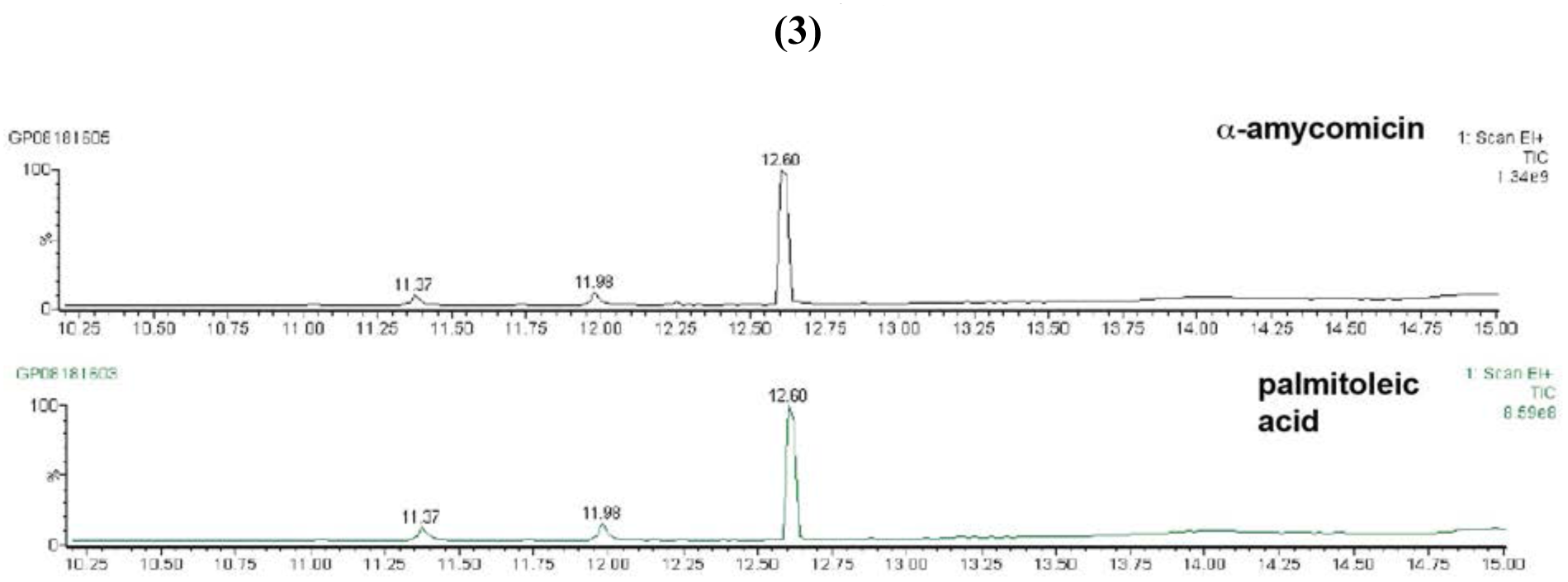
GCMS chromatogram of naturally-produced and authentic palmitoleic acid (3)

**Fig. S7:**
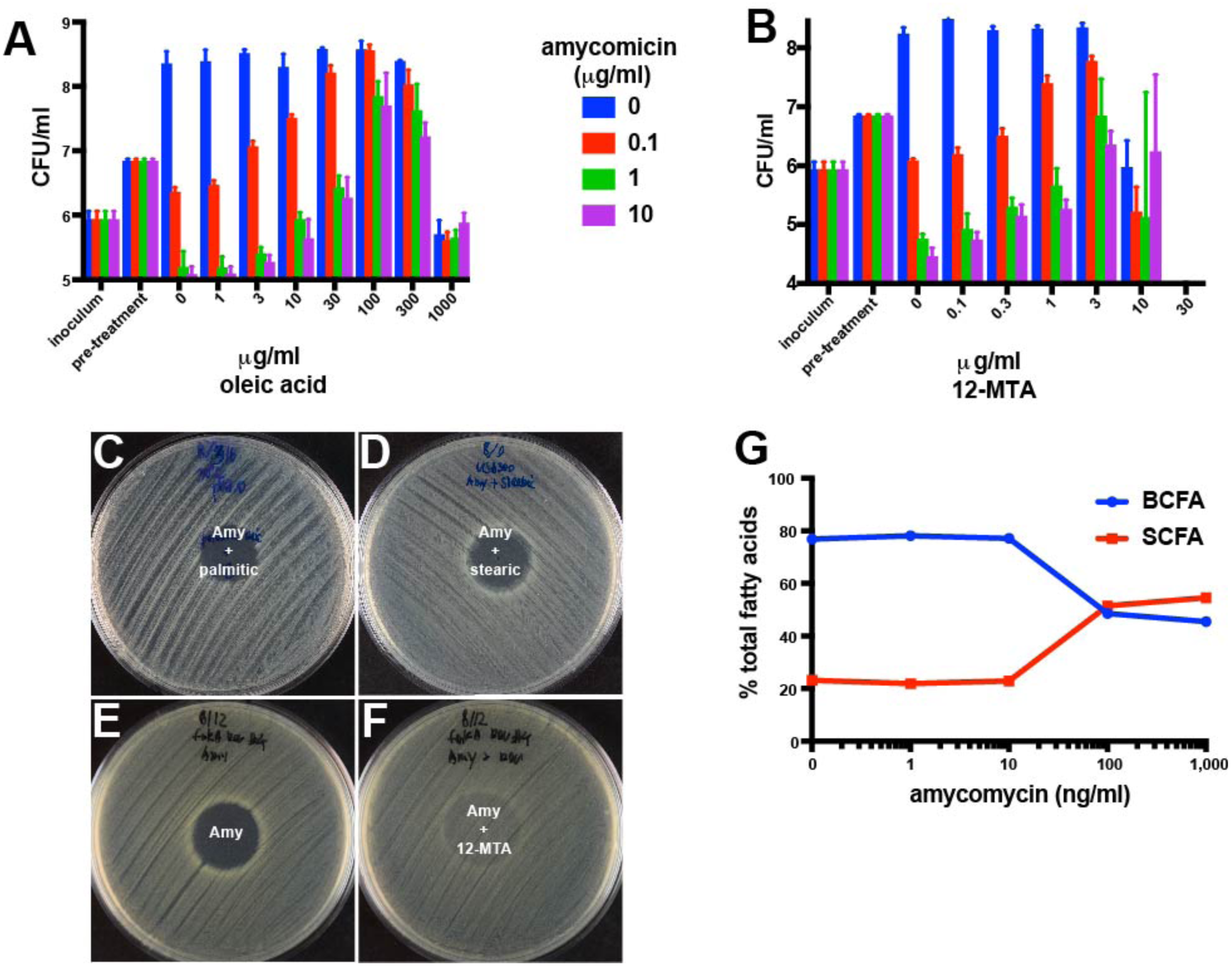
The effect of fatty acids on amycomicin activity against *S. aureus*. (*A* and *B*) CFUs of *S. aureus* detected upon treatment with (*A*) amycomicin plus the indicated concentrations of oleic acid and (*B*) amycomicin plus 12-methyltetradecanoic acid* (limit of detection was103 CFU/ml). (*C* and *D*) Zones of inhibition on lawns of *S. aureus* produced by amycomicin with palmitic or stearic acid. Isolates of *fakA* suppressor mutants that proliferated in the presence of amycomicin and 12-MTA (Fig. 3F) unable to proliferate in: (*E*) pure amycomicin, but growing when (*F*) 12-MTA is supplemented with amycomicin. (*G*) Fatty acid profile of *S. aureus* treated with indicated concentrations of amycomicin. Error bars indicate standard deviation. *No live bacteria were detected at or above 30 *μ*g/ml 12-MTA

**Fig. S8:**
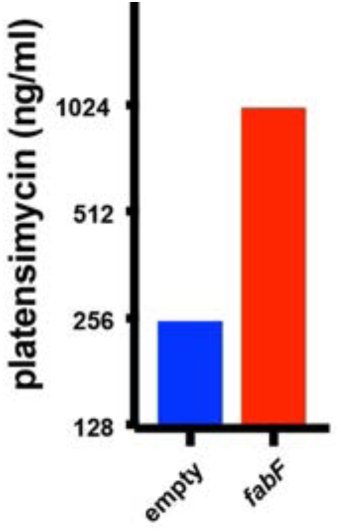
Minimal inhibitory concentration of platensimycin against *S. aureus* containing an empty plasmid or a plasmid overexpressing *fabF*.

**Fig. S9:**
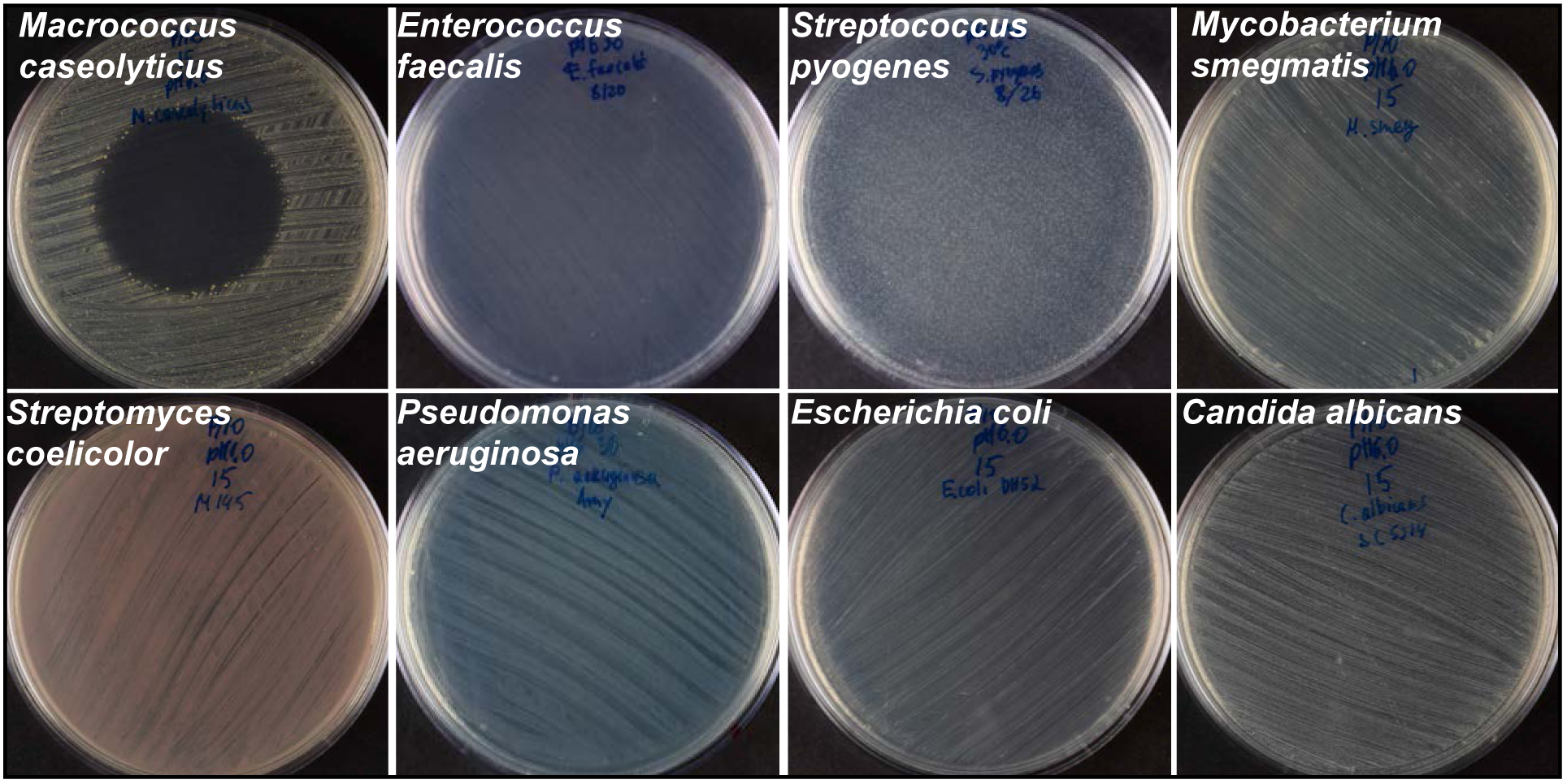
Various microbial species treated with amycomicin. Effect of adding 1 *μ*g amycomicin to lawns of the indicated microorganisms.

**Fig. S10:**
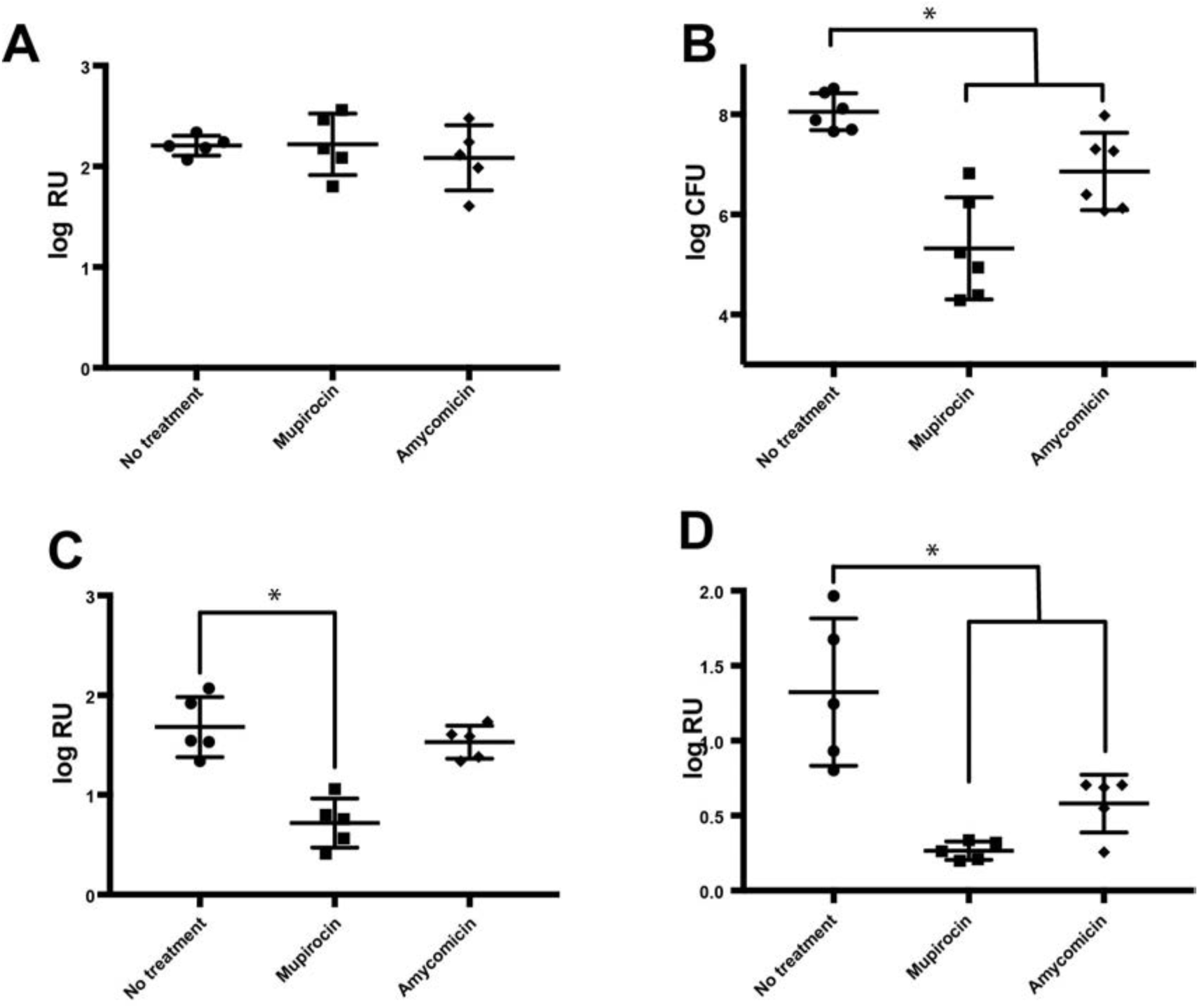
Amycomicin treatment of mice infected with *S. aureus*. (*A*) Bioluminescence expressed as radiance units (RU) detected in the infected sites 1-day post-infection, before any treatment was applied. (*B*) Colony forming units (CFU) isolated from the infected sites of mice infected with *S. aureus* after three daily doses of indicated treatments. (*C* and *D*). Bioluminescence expressed in radiance units (RU) detected within the infected sites after (*C*) one or (*D*) two days of the indicated treatments. Horizontal bars indicate averages and standard deviations of log_10_-transformed values. Welch’s t-test was used to determine the p-values. P-values that are smaller than 0.05 are marked with an asterisk.

**Fig. S11:**
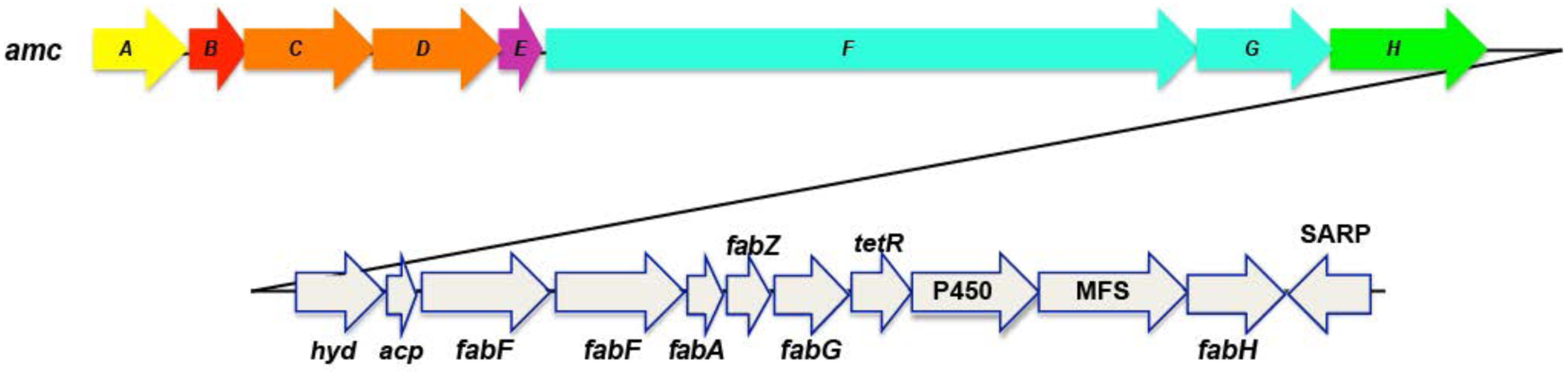
Genes downstream from *amcA-H* gene cluster. Putative functions: *hyd* – α/β hydrolase; *acp* – acyl-carrier protein; *fabF -* 3-oxoacyl-[acyl-carrier-protein] synthase 2, KASII; *fabA* - 3-hydroxydecanoyl-[acyl-carrier-protein] dehydratase; *fabZ -* 3-hydroxyacyl-[acyl-carrier-protein] dehydratase; *fabG* - 3-oxoacyl-[acyl-carrier protein] reductase; *tetR* –TetR family transcriptional regulator; P450 – cytochrome P450; MFS – major facilitator superfamily transmembrane transporter; *fabH* - 3-oxoacyl-[acyl-carrier-protein] synthase, KASIII; SARP – SARP family transcriptional regulator.

**Table S3.** Percent fatty acid composition of *S. aureus* cells treated with amycomicin.

| fatty acid | Amycomycin concentration |  |  |  |  |
| --- | --- | --- | --- | --- | --- |
|  | 0 ng/ml | 0.1 ng/ml | 1 ng/ml | 10 ng/ml | 100 ng/ml |
| **12:0** | 0.08 | 0.09 | 0.10 | 0.61 | 0.74 |
| **13:0 iso** | 0.22 | 0.23 | 0.24 | 0.00 | 0.00 |
| **13:0 anteiso** | 0.07 | 0.07 | 0.07 | 0.00 | 0.00 |
| **14:0 iso** | 7.51 | 7.94 | 8.05 | 5.94 | 6.08 |
| **14:0** | 0.77 | 0.76 | 0.75 | 1.74 | 1.96 |
| **15:1 iso G** | 0.05 | 0.05 | 0.07 | 0.00 | 0.00 |
| **15:0 iso** | 12.10 | 12.27 | 11.79 | 4.70 | 4.65 |
| **15:0 anteiso** | 37.28 | 37.99 | 37.26 | 25.19 | 22.10 |
| **15:0** | 0.09 | 0.08 | 0.08 | 0.00 | 0.00 |
| **16:0 iso** | 5.37 | 5.59 | 5.43 | 4.03 | 4.14 |
| **16:0** | 3.49 | 3.40 | 2.98 | 5.37 | 5.52 |
| **15:0 2OH** | 0.00 | 0.00 | 0.04 | 0.00 | 0.00 |
| **17:0 iso** | 4.15 | 4.10 | 3.97 | 1.23 | 1.27 |
| **17:0 anteiso** | 6.65 | 6.51 | 6.49 | 3.86 | 3.42 |
| **17:0** | 0.34 | 0.34 | 0.32 | 0.00 | 0.00 |
| **18:0 iso** | 1.58 | 1.58 | 1.69 | 1.41 | 1.47 |
| **18:2 w6,9c** | 0.13 | 0.12 | 0.13 | 0.91 | 0.97 |
| **18:1 w9c** | 0.10 | 0.07 | 0.09 | 0.59 | 0.75 |
| **18:1 w7c** | 0.06 | 0.07 | 0.07 | 0.52 | 0.61 |
| **18:0** | 12.00 | 11.34 | 11.17 | 17.11 | 17.06 |
| **19:0 iso** | 0.87 | 0.83 | 0.93 | 0.53 | 0.59 |
| **19:0 anteiso** | 0.81 | 0.77 | 0.87 | 0.80 | 0.76 |
| **19:0** | 0.58 | 0.56 | 0.63 | 0.92 | 0.97 |
| **18:0 3OH** | 0.05 | 0.05 | 0.09 | 0.00 | 0.00 |
| **20:0 iso** | 0.22 | 0.21 | 0.32 | 0.85 | 0.98 |
| **20:2 w6,9c** | 0.06 | 0.05 | 0.04 | 0.98 | 0.00 |
| **20:0** | 5.35 | 4.92 | 6.36 | 22.73 | 25.96 |

**Table S4.**
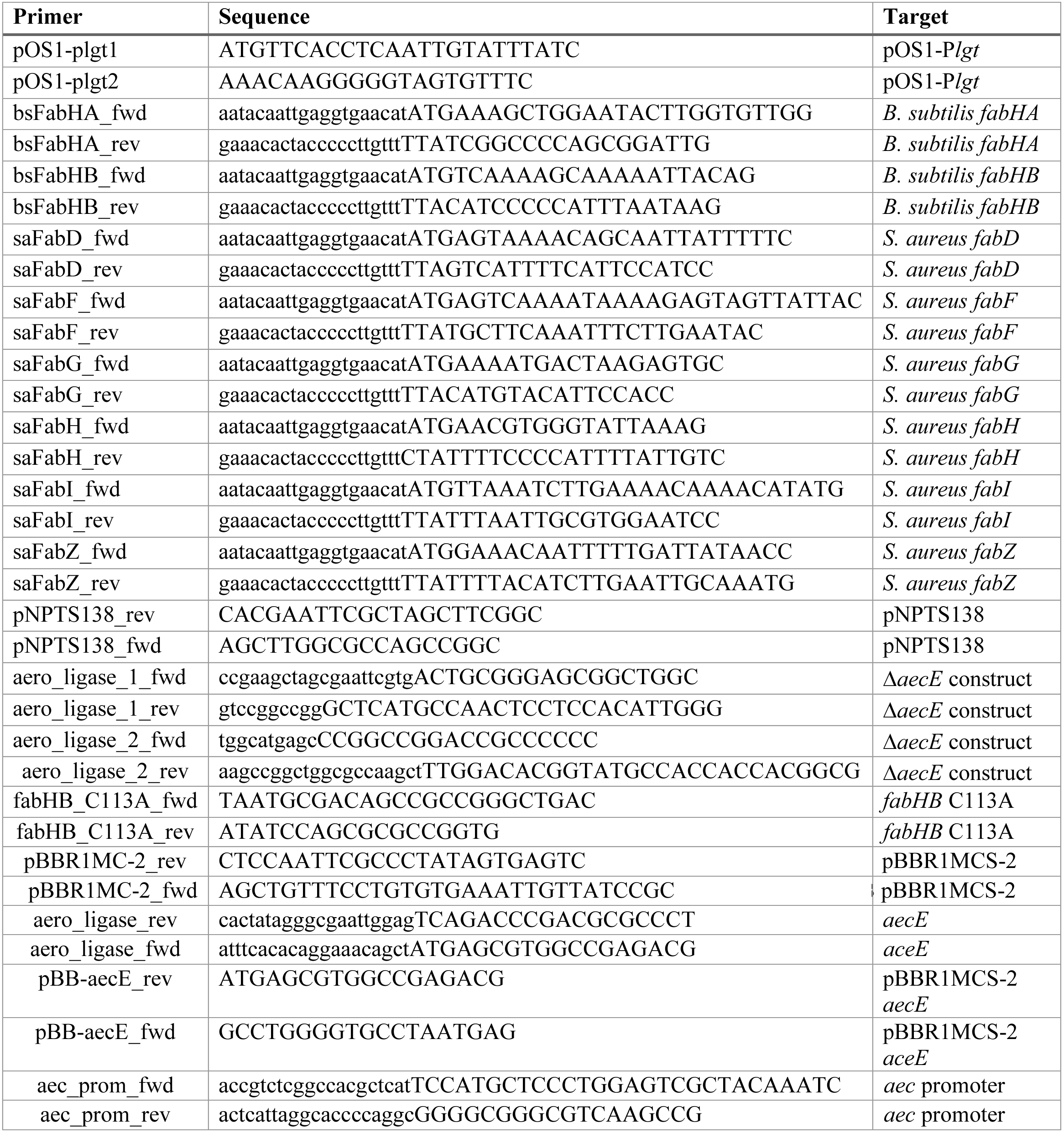
Primers used in the study.

